# Physiological causes and biogeographic consequences of thermal optima in the hypoxia tolerance of marine ectotherms

**DOI:** 10.1101/2022.02.03.478967

**Authors:** Martin-Georg A. Endress, Thomas H. Boag, Benjamin P. Burford, Justin L. Penn, Erik A. Sperling, Curtis A. Deutsch

## Abstract

The minimum O_2_ needed to fuel the demand of aquatic animals is commonly observed to increase with temperature, driven by accelerating metabolism. However, recent measurements of critical O_2_ thresholds (‘*P*_*crit*_’) reveal more complex patterns, including those with a minimum at an intermediate thermal ‘optimum’. To discern the prevalence, physiological drivers, and biogeographic manifestations of such curves, we analyze new experimental and biogeographic data using a general dynamic model of aquatic water breathers. The model simulates the transfer of oxygen from ambient water, through a boundary layer and into animal tissues driven by temperature-dependent rates of metabolism, diffusive gas exchange, and ventilatory and circulatory systems with O_2_-protein binding. We find that a thermal optimum in *P*_*crit*_ can arise even when all physiological rates increase steadily with temperature. This occurs when O_2_ supply at low temperatures is limited by a process that is more temperature sensitive than metabolism, but becomes limited by a less sensitive process at warmer temperatures. Analysis of species respiratory traits suggests this scenario is not uncommon in marine biota, with ventilation and circulation limiting supply under cold conditions and diffusion limiting supply at high temperatures. Using biogeographic data, we show that species with these physiological traits inhabit lowest O_2_ waters near the optimal temperature for hypoxia tolerance, and are restricted to higher O_2_ at temperatures above and below this optimum. Our results imply that O_2_ tolerance can decline under both cold and warm conditions, and thus may influence both poleward and equatorward species range limits.

**Significance Statement:** Physiology shapes the ecology, biogeography, and climate responses of marine species. In aquatic ectotherms, accelerating metabolism and lowered oxygen availability generally result in increasing oxygen limitation with warming. Here we present evidence for thermal optima in hypoxia tolerance of diverse species that is explained by a dynamical model of organismal physiology. Our results indicate that this potentially widespread bidirectional pattern explains species biogeographic limits in cold and warm waters. It can be understood using a generalized Metabolic Index of O_2_ supply to demand, which captures the variable observed trends between temperature and species hypoxia sensitivity. Oxygen limitation of aerobic metabolism in cold water has far-reaching implications for marine biogeography and species migrations under climate change.

## Introduction

Climate change is raising temperatures throughout the upper ocean, while decreasing its oxygen content. These trends are among the most robustly observed and well understood aspects of global ocean change (1). They also pose a major challenge for marine ectotherms, whose metabolic rates rise exponentially with temperature (2,3), requiring a concomitant increase in O_2_ supply to maintain aerobic energy balance that is at odds with the ocean’s declining global O2 inventory (4, 5). The temperature-dependent hypoxia tolerance of marine species already limits their geographic distributions, most commonly at the equatorial (warm) and/or deep (low O_2_) range edge of species distributions (6, 7, 8, 9), yielding a simple physiological mechanism for species responses to climate change (10, 11).

The environmental O_2_ minimum at which an organism can sustain its resting metabolism is typically reported as a critical pressure (*P*_*crit*_) and remains the most common measure of hypoxia tolerance, despite potential complexities of experimental determination (12, 13). In most studied species, *P*_*crit*_ increases with temperature, implying that their O_2_ demand accelerates faster with warming than their supply (6).

Some species show a decrease in *P*_*crit*_ as temperatures rise, implying that supply accelerates faster than demand, although this has rarely been observed (14, 15). In recent experiments, still other species exhibit both a decline in *P*_*crit*_ as temperatures rise from the coldest water, followed by an increase from further warming, resulting a minimum Pcrit, and thus a maximum hypoxia tolerance, at an intermediate optimum temperature (16, 17). While the individual processes of supply and demand all tend to increase steadily with temperature (18, 19), these bowl-shaped *P*_*crit*_ curves require that the ratio of these rates exhibits a more complex relationship to temperature. Thermal optima for hypoxia tolerance have been posited (20, 21), but scarce empirical support has hampered the development of a quantitative model. This prevents a mechanistic evaluation of the role of hypoxia tolerance at the cold edge of range limits, and the associated implications of climate change, especially for populations not living near a species’ warm range limit, or exposed to ocean cooling.

To examine the prevalence of thermal optima in hypoxia tolerance, diagnose the physiological conditions under which it can arise, and evaluate its relevance to species biogeography, we combined new laboratory experiments, a dynamic model of O_2_ supply in marine ectotherms, and species biogeographic distribution data. Among all studied species, we find complex *P*_*crit*_ behavior across a broad temperature range. A general model of aquatic water breathers demonstrates the conditions under which thermal optima can emerge from the multi-step nature of the O_2_ supply chain, and analysis of prior laboratory data suggests that marine species commonly meet those conditions. The behavior of the dynamic model can be reproduced with a generalized Metabolic Index of O_2_ supply to demand (6) that captures a wide range of observed *P*_*crit*_ curves. Finally, we present evidence that *P*_*crit*_ curves with a thermal optimum are also reflected in the biogeography of marine species and thus may explain the cold/poleward limit in such species geographic ranges.

## Results

### Laboratory Observations

To evaluate the prevalence of non-exponential *P*_*crit*_ curves, we measured *P*_*crit*_ across a wide temperature spectrum for four previously unmeasured invertebrate species. These species span four different phyla, have multiple modes of oxygen supply, and regularly encounter temporal and/or spatial variability in environmental pO_2_ (Fig. 1). Following published respirometry protocols (Materials and Methods), we conducted *P*_*crit*_ measurements from the freshwater oligochaete worm *Tubifex tubifex*, the outer shelf/upper slope sea urchin *Lytechinus pictus* from California, and the Atlantic intertidal anemone *Nematostella vectensis*. We also used *P*_*crit*_ measurements of the squid D*oryteuthis opalescens* which is exposed to strong gradients of temperature and O_2_ in the California Current System (22).

**Fig. 1.**
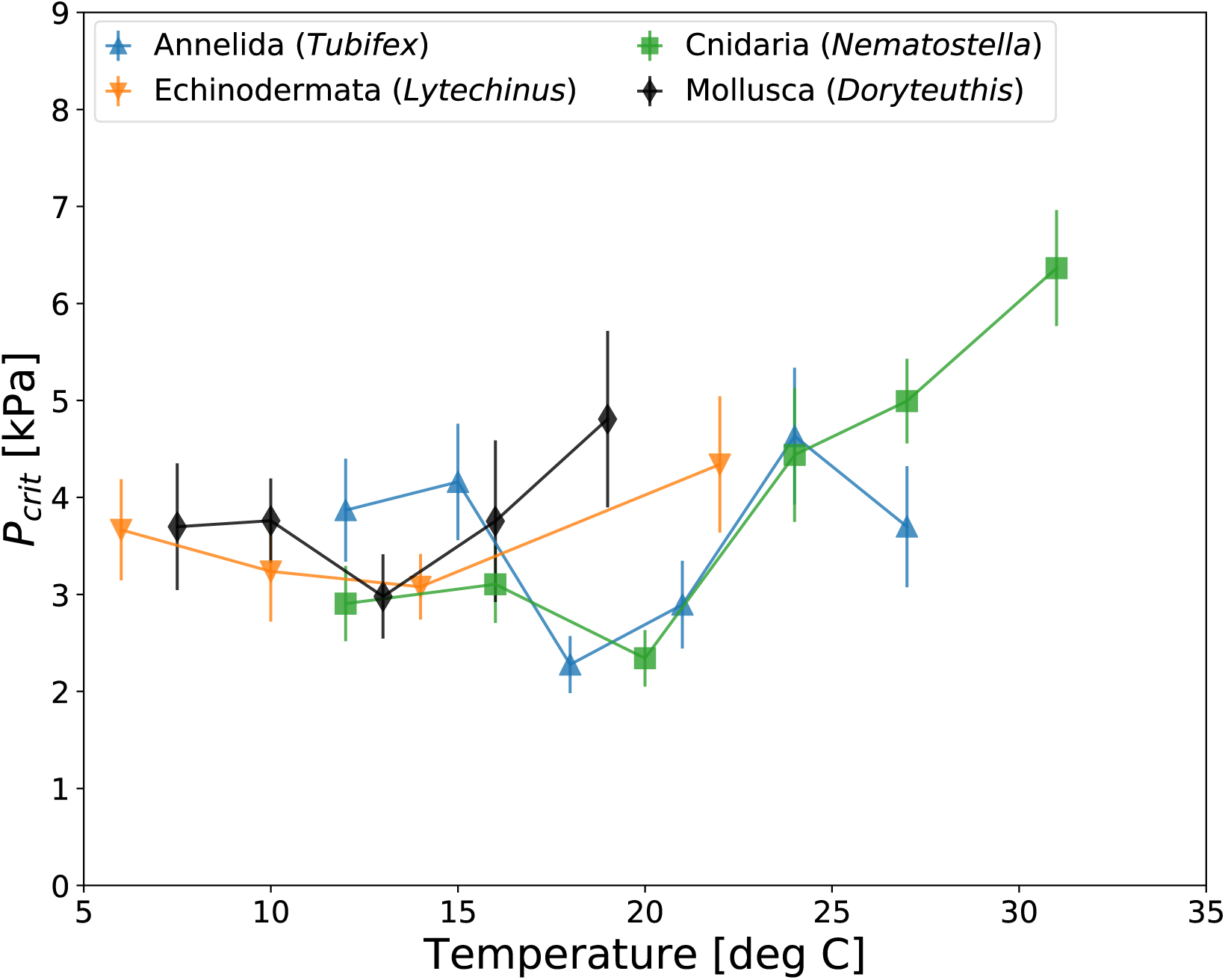
Temperature-dependent critical oxygen pressures (*P*_*crit*_, mean ± SE) of 4 marine invertebrate species from new closed-system respirometry experiments exhibit a minimum at intermediate temperatures, indicating a thermal optimum in hypoxia tolerance. The species include an oligochaete worm (*Tubifex tubifex*), a sea urchin (*Lytechinus pictus*), an anemone (*Nematostella vectensis*) and a cephalopod (*Doryteuthis opalescens*). See text for details.

Measurements of all species reveal a minimum in *P*_*crit*_ at intermediate experimental temperatures, with substantial variation in the location of the thermal optimum and depth of the *P*_*crit*_ minimum. Some species *P*_*crit*_ curves reveal a broad bowl (*L. pictus*), others a deep bowl (*T. tubifex*), and still others a relatively constant *P*_*crit*_ at cold temperatures, followed by a sharp rise at warmer temperatures (*N. vectensis, D. opalescens*). These new respirometry data combined with published data (17,16), indicate that thermal optima in hypoxia tolerance are found in multiple phyla and across multiple modes of oxygen supply (e.g., gills and a blood vascular system in squid versus cutaneous respiration in anemones) and may therefore represent a widespread pattern.

### Dynamic Model

To explore the conditions that lead to a thermal optimum in hypoxia tolerance, we develop a dynamic model of O_2_ supply and demand in water-breathing animals. The model simulates the transfer of O_2_ from the environment to the metabolizing tissues of an organism across a range of temperatures using a system of coupled non-linear ordinary differential equations (ODEs). To make the model generally applicable to aquatic animals, we include all the potential pools and fluxes of O_2_, including external ventilation of water from the ambient fluid to the boundary layer at the exchange surface, the molecular O_2_ diffusion across that surface, and internal flux of O_2_ to metabolizing body tissues, which may be mediated by a circulatory system (Fig. 2A). In addition to dissolved O_2_, the model also tracks the concentration of the bound and unbound forms of an oxygen-transport protein such as hemoglobin or hemocyanin (denoted HxO and Hx, respectively), which bind and release molecular O_2_ according to the associated chemical equilibrium. This is captured by the pO_2_ at half-saturation (denoted *P*_50_) and the enthalpy (Δ*H*) of the binding reaction, which governs the temperature dependence of that equilibrium (23, details SI).

**Fig. 2.**
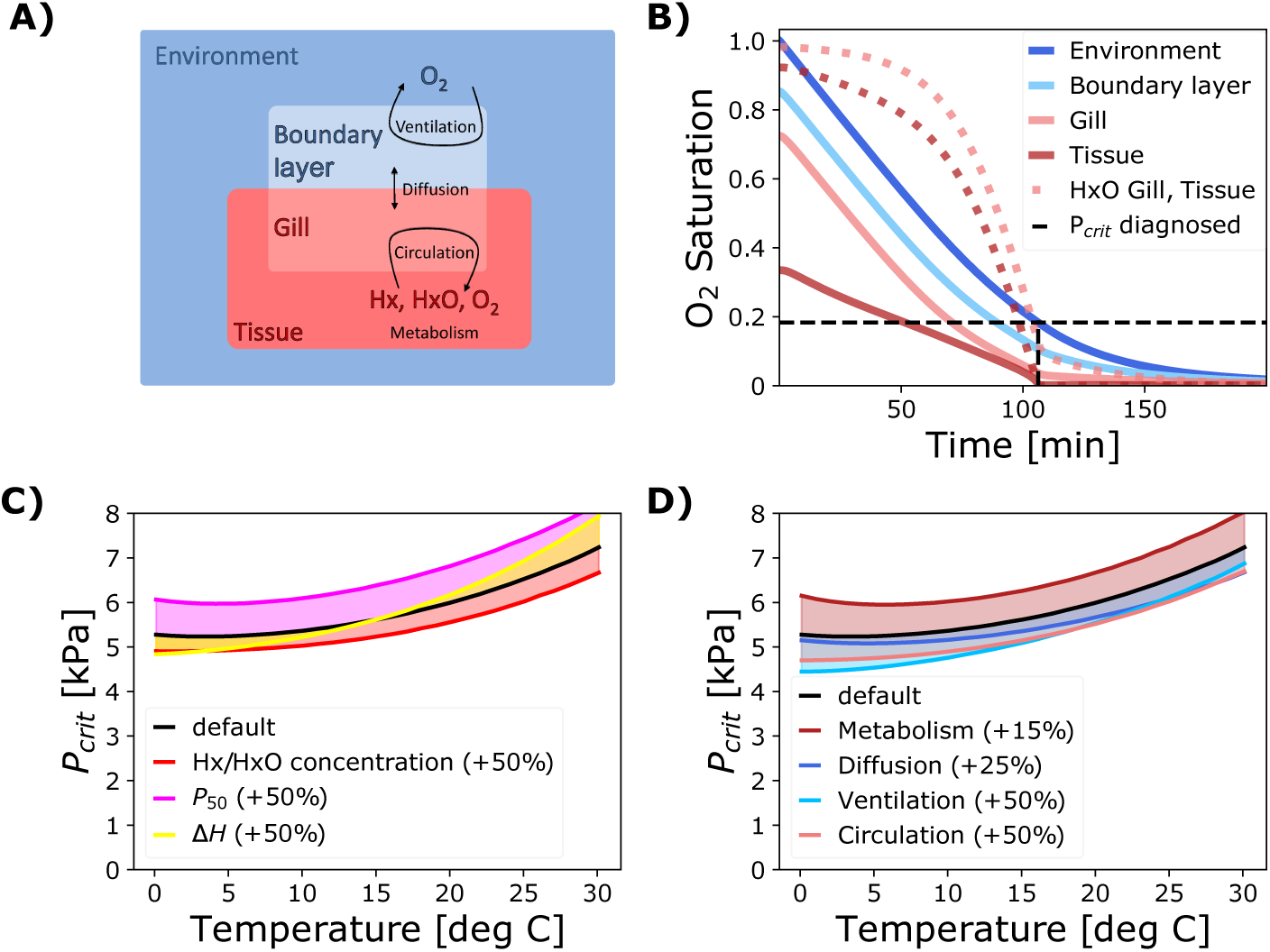
Structure and output of the dynamical model used to investigate the effects of a multistep O_2_ supply chain. **A)** The model tracks the concentrations of O_2_ as well as unbound and bound O_2_ transporting proteins (’Hx’, ‘HxO’) in 4 compartments representing external and internal volumes of water or body fluid, which are connected through a linear O_2_ supply chain with external ventilation, diffusion and internal circulation. **B)** A model run at a single temperature resembles a closed system respirometry experiment. The saturation of O_2_ (solid colored) and proportion of HxO (dashed colored) decline in all compartments until the O_2_ level in the metabolizing tissue (dark red) reaches a critical limit near zero, at which point metabolic consumption slows down and *P*_*crit*_ can be determined from the rate of environmental O_2_ depletion (dashed black). **C)** Effects of increasing the concentration, half-saturation pressure (*P*_50_) or temperature sensitivity (Δ*H*) of O_2_ transport protein on the *P*_*crit*_ curve. **D)** Effects of increasing the rate coefficients of biophysical supply and demand processes. A higher metabolism elevates the curve, while increasing the rate of any supply process lowers it.

Each of the three O_2_ supply processes (ventilation, diffusion, and circulation) is described by a rate *S*_*i*_ that is represented as the product of the pO_2_ difference between the respective compartments, Δ_*i*_ *pO*_2_, and a temperature-dependent rate coefficient 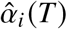 that characterizes the kinetics of that process:

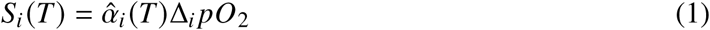

The temperature-dependency of the three rate coefficients – flow rates of ventilated water and circulated blood, and the diffusivity of O_2_ – each vary exponentially with temperature (Arrhenius function), as does the metabolic rate, but with distinct temperature sensitivities. The resulting 8 parameters (3 supply rate coefficients, the metabolic rate, and the temperature sensitivity of each) along with the 3 chemical parameters (*P*_50_, Δ*H* and total Hx concentration) represent a set of traits that determine a model organism’s hypoxia tolerance and its variation with temperature. The well-documented trait variations in real animals (e.g., overall O_2_ supply capacity [24] and adaptation to hypoxia [25], gill surface area [26, 27], blood properties [28, 29]) are simulated by scaling these parameters in the model. Our analysis aims to discern how such biological traits govern the shape of the resulting *P*_*crit*_ curves with respect to temperature.

Model simulations resemble standard closed system respirometry experiments used to determine *P*_*crit*_ values (Fig. 2B, 30), in which O_2_ is depleted from the ambient water as it gets transferred to metabolizing tissues. Both the O_2_ concentrations and the fraction of O_2_-bound protein (HxO) decline in all compartments. Once O_2_ levels in the tissue compartment can no longer support resting metabolism, consumption slows down with the onset of hypoxemia, allowing *P*_*crit*_ to be diagnosed from the rate of environmental O_2_ depletion using breakpoint analysis (Materials and Methods, full model in SI).

Simulations across a range of temperatures yield the *P*_*crit*_ curve, which integrates the contribution of all traits to a single metric of hypoxia tolerance. Across a wide range of model parameters centered on the most common traits observed in marine organisms (7), the *P*_*crit*_ curves exhibit an overall rise with temperature, driven by the increase in metabolic rate.

Both the chemical properties (Hx, Δ*H* and p50) as well as the rate coefficients of supply and demand 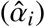 have intuitive impacts on the *P*_*crit*_ curves. For example, a higher concentration of total Hx acts to lower the *P*_*crit*_ curve across all temperatures (Fig. 2C), enhancing the tolerance to hypoxia. An equivalent effect can be obtained by increasing the biophysical supply coefficients, simulating changes such as a larger gill area or faster ventilation rate (Fig. 2D).

However, we also find less intuitive impacts on the shape of the curves, as the fractional change in *P*_*crit*_ is not always the same across the full temperature range. For instance, a 25 % increase in the ventilation rate does not lower the *P*_*crit*_ by the same fraction at all temperatures, but instead has a larger impact under cold conditions than under warm conditions (Fig. 2D). In other words, the *P*_*crit*_ curves resulting from a multi-step supply chain can depart from simple exponential relationships with temperature, even when each single supply process accelerates exponentially with warming. We conclude that the well-known non-linearities in blood-O_2_ binding are not the essential cause of this behavior, because the variation due to biophysical properties is similar to that induced by variations in blood chemistry. Moreover, we observe complex *P*_*crit*_ curves in organisms without O_2_-binding proteins (e.g. *N. vectensis* in Fig. 1).

Instead, we focus our analysis on the mechanisms by which the linear combination of biophysical transfer processes in a multi-step O_2_ supply chain leads to the complex patterns observed in *P*_*crit*_ curves.

The origins of non-exponential *P*_*crit*_ curves can be demonstrated quantitatively in a model with a supply chain consisting only of ventilation and diffusive gas exchange (Fig. 3). In isolation, each step yields a simple (exponential) *P*_*crit*_ curve with a slope depending on the temperature sensitivities of supply and demand. The curve is increasing if metabolic demand accelerates faster with temperature than supply (shown for diffusion, Fig. 3A), and decreasing if instead the temperature sensitivity of supply exceeds that of metabolism (ventilation, Fig. 3B). Combining ventilation and diffusion in series results in a *P*_*crit*_ curve that is the sum of the two curves corresponding to the single steps, and thus exhibits a minimum at an intermediate temperature (Fig. 3C).

**Fig. 3.**
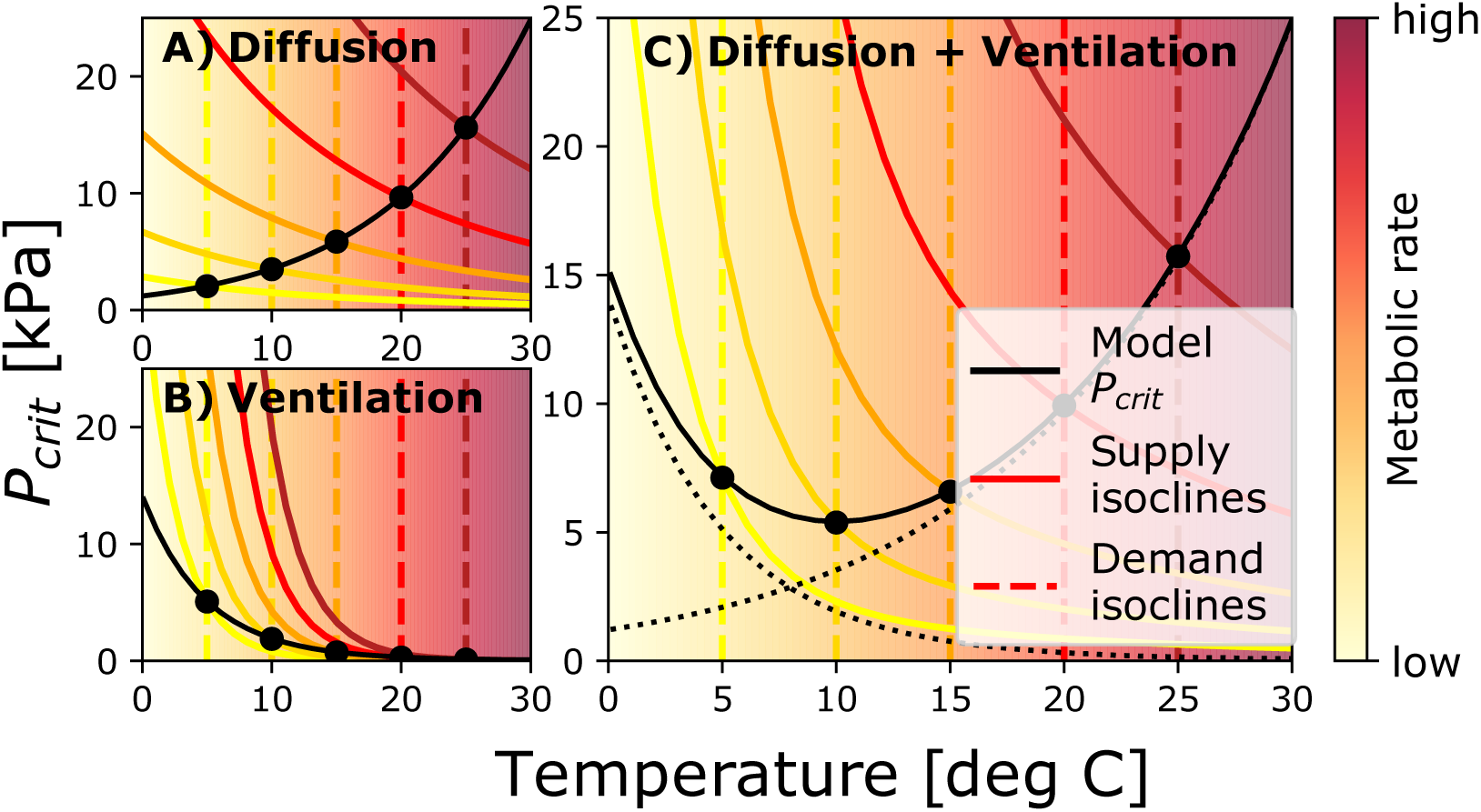
Quantitative analysis of the *P*_*crit*_ curve of a model with a two-step supply chain consisting of ventilation and diffusive gas exchange. At any given temperature, the model *P*_*crit*_ can be found analytically as the intersection (black dots) of the demand isocline (dashed colored) and the corresponding supply isocline (solid colored), aligning with the curve diagnosed from numerical simulations (solid black). **A)** In a model with only diffusive gas exchange characterized by a smaller temperature sensitivity than metabolic demand, the isocline intersections yield an increasing *P*_*crit*_ curve. **B)** Conversely, the steeper supply isoclines lead to a decreasing pattern in a single supply step model with ventilation that accelerates faster than metabolic demand. **C)** Combining the two supply steps in series results in a *P*_*crit*_ curve that is the sum of the single step curves (dotted black), giving rise to a thermal optimum at intermediate temperatures.

This additive nature of the *P*_*crit*_ curve resulting from a linear supply chain can also be derived analytically from the system of model ODEs for more than two supply steps (details in SI). Conceptually, this property can be thought of as analogous to an electrical circuit in which a fixed voltage is applied to a series of resistors. Just like the total voltage can be obtained as the sum of the individual voltage drops across each resistor, the total *P*_*crit*_ curve of a multi-step supply chain can be obtained as the sum of the pO_2_ drops that drive each individual supply process. Therefore, a bowl-shaped *P*_*crit*_ curve can emerge if the supply chain includes processes that are both more and less sensitive to temperature changes than metabolism. In Fig. 3C, the *P*_*crit*_ curve rises under warm conditions because a large pO_2_ gradient is required to drive sufficient diffusion at high temperatures. This is due to the fact that diffusion accelerates slower than metabolism with warming. On the other hand, the curve also remains flat or even reverses under cold conditions because a large pO_2_ gradient is required to drive sufficient ventilation at low temperatures, since this process has a higher temperature sensitivity than metabolism.

Because the critical pO_2_ differences required to drive the individual supply steps are not the same, the total *P*_*crit*_ curve is not equally sensitive to changes in the biologically controlled rate coefficients at all temperatures. In the example above, the change in *P*_*crit*_ at high temperatures due to a change in ventilation rate might be small or even negligible while its response to a change in diffusivity might be substantial, even for the same relative increase in the biologically controlled parameter. More generally, a change in the coefficient of any supply process that accelerates faster with warming than metabolism will have the largest impact on *P*_*crit*_ under cold conditions, as in the case of ventilation. On the other hand, such an increase has the largest impact on *P*_*crit*_ under warm conditions for a supply process that accelerates slower than metabolism, such as diffusion.

This relationship is particularly important for processes under immediate biological control like ventilation and circulation and has implications for understanding their temperature sensitivity. Incurring the energetic costs of accelerating heart rate or ventilation across the entire temperature range may not be beneficial if *P*_*crit*_ is instead much more sensitive to changes in diffusion at high temperatures. We illustrate this in a model variant with a ventilation rate that has a high temperature sensitivity at low temperatures but reaches an upper limit under warm conditions, as for example observed in (31, 32, 33). The resulting change in *P*_*crit*_ at high temperatures compared to a simple exponential ventilation rate is minimal (Fig. S3). In this scenario, increasing the ventilation rate throughout the warm side of the temperature range barely impacts hypoxia tolerance, because O_2_ supply is largely determined by diffusion.

### Evidence from Physiology

To determine whether the physiological conditions for a thermal optimum in hypoxia tolerance are common among marine biota, we compiled experimental data on the temperature dependence of ventilation and circulation rates of aquatic water breathers (Materials and Methods).

The compilation covers 58 data sets from 35 species, including 21 chordates, 9 arthropods, 3 annelids and 2 mollusks. Estimates of the sensitivity parameters (*E*_*V*_, *E*_*C*_) obtained by fitting Arrhenius functions to the data show an increase in ventilatory and circulatory activity with temperature in almost all species, with sensitivities ranging from −0.14 eV to 0.9 eV and a mean of 0.39 eV (± 0.22 eV SD). Since these estimates include frequencies and stroke volumes in addition to volumetric flow rates, they represent a lower bound on the actual sensitivity of ventilation and circulation as considered in the dynamic model. A higher sensitivity (0.49 eV ± 0.21 eV SD) is obtained if only volumetric rates (n =12) are considered (Fig. 4).

**Fig. 4.**
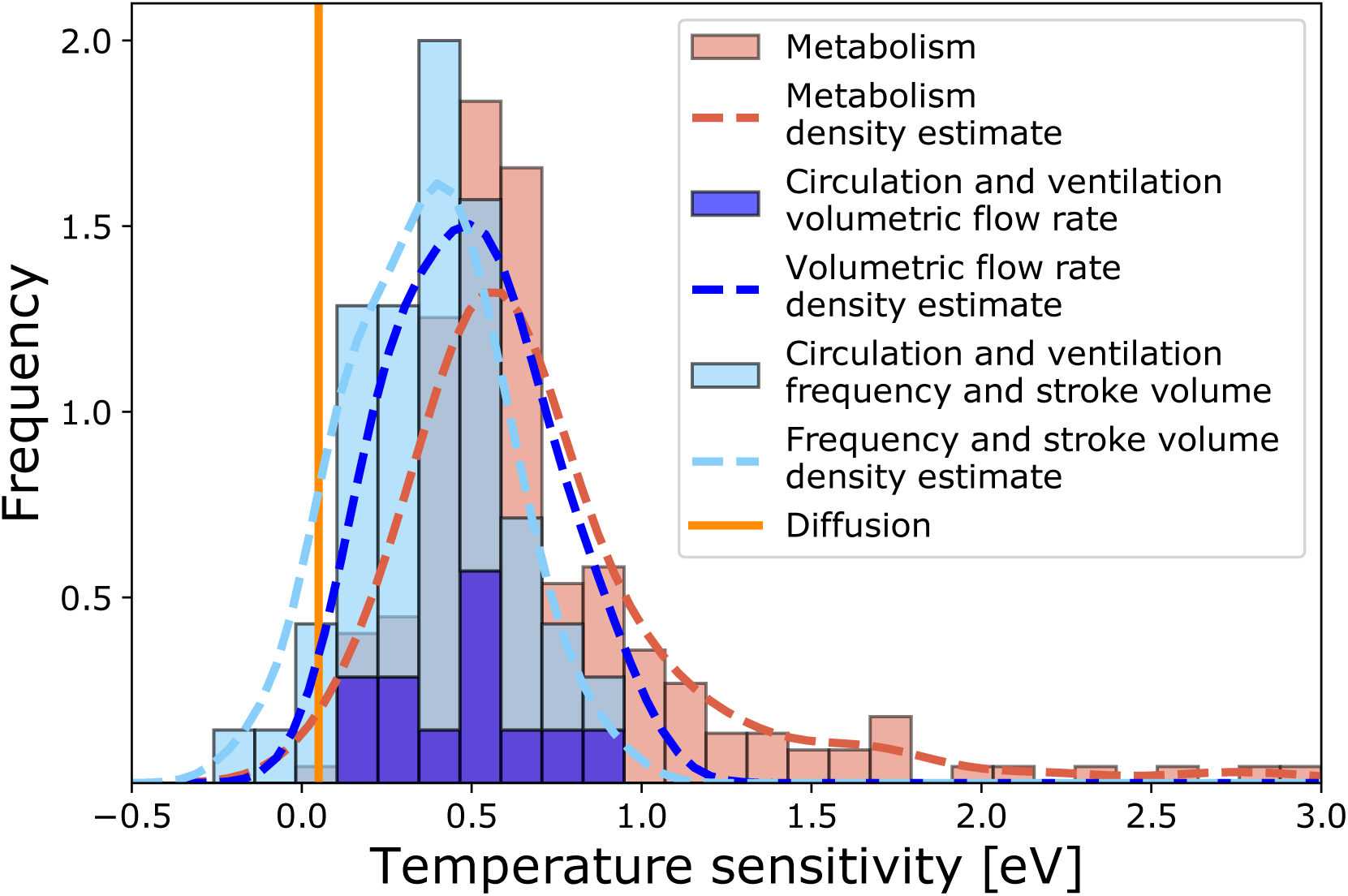
The temperature sensitivities of ventilation and circulation rates estimated from published experimental data (blue, 58 estimates from 35 species) fall between the theoretical prediction for the sensitivity of diffusion (vertical orange line) and existing estimates for the sensitivity of metabolic rates (186 species, [7]) on average. Dashed lines show kernel density estimates of the trait frequency distributions.

These results can be compared to existing estimates of the temperature sensitivity of metabolism (*E*_*M*_) with a mean of 0.71 eV [± 0.46 eV SD] from a diverse set of 186 species (7). If the traits of O_2_ supply and demand were independent, the estimated frequency distributions in Fig. 4 would suggest that the conditions for thermal optima (*E*_*V*_, *E*_*C*_ > *E*_*M*_) are met in about 23 % of species after accounting for the effect of decreasing solubility with temperature (SI). However, supply sensitivities exceed that of demand in 7 of 17 species for which both estimates are available (Dataset S1). Thus, about 40 % of species with adequate data meet this condition for having a thermal optimum in hypoxia tolerance.

The thermal variation of ventilation and circulation rates also differs across the inhabited temperature range for many species. We estimated the temperature sensitivity of volumetric flow rates in both warm and cold temperature ranges, for all species with sufficient data (*n* = 10). On the cold side the mean *E*_*V*_ and *E*_*C*_ are 0.69 eV, significantly exceeding the cold side with a mean of 0.07 eV (p = 0.009). The difference remains significant if stroke volumes and frequencies are also considered (*n* = 40 from 25 species, Fig. S5, details in SI). Thus, the acceleration of ventilation and circulation rates slows down in the warmer half of experimental temperatures on average. This behavior is consistent with these biophysical processes conferring little additional hypoxia tolerance at high temperatures for the associated energetic cost, with *P*_*crit*_ being most sensitive to diffusion as illustrated in the model variant for ventilation (Fig. S3). Taken together, both the high temperature sensitivity of biophysical rates (circulation and ventilation) in colder waters, and the reduction of these sensitivities in warmer waters, suggest that the condition for thermal optima are commonly found among the traits of marine species.

### A Metabolic Index with Thermal Optima

We generalized the Metabolic Index introduced by Deutsch et al. (6) to account for the occurrence of complex shaped *P*_*crit*_ curves. The index is defined as the ratio of O_2_ supply to resting demand of an aquatic water-breather in its environment, and it has been applied to understand how species biogeography is shaped by climate (6, 16, 11, 10, 34). However, the original formulation assumed that *P*_*crit*_ varies exponentially with temperature. Our generalized version is able to reproduce the full range of behaviors exhibited by the dynamic model, requiring only five parameters, which can be calibrated from experimental data through a single equation.

While it is possible to derive the ratio of supply to demand from the model ODEs analytically (SI), the metabolic index can also be the developed from the analogy of an electrical circuit in which a fixed voltage is applied to a series of resistors.

When considering a single supply step *i*, the rate of O_2_ supply according to Eqn. (1) is the product of the pO_2_ difference Δ*pO*_2_ between the compartments, equivalent to a ‘voltage’ driving a current, and the biologically determined rate coefficient 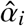 of the process, equivalent to a temperature-dependent ‘conductance’ with respect to the flow of O_2_.

For the habitat to be viable, this supply rate must be equal to metabolic consumption, such that *P*_*crit*_ can be interpreted as the minimum ‘voltage’ required to achieve an O_2_ supply matching demand given the fixed ‘conductance’ of the biological supply process.

In a supply chain with multiple steps in series, each step is associated with such a required voltage drop - a pO_2_ difference - determined by its single step conductance. Thus, the *P*_*crit*_ of the composite chain can be obtained as the sum of the minimum pO_2_ differences of the single supply steps, as illustrated in Fig. 3C.

The temperature-dependent ‘conductance’ (or rate coefficient) 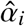 of a single supply step can be expressed as 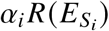, where *α*_*i*_ denotes the value of the coefficient at reference temperature, which is scaled by an exponential (Arrhenius) function *R* with temperature sensitivity 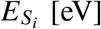 [eV]. More generally, in a chain with *n* supply steps in series, the total conductance of the chain is the reciprocal of the sum of single step resistances. When divided by metabolic demand *α*_*M*_ *R*(*T, E*_*M*_), the resulting expression for the generalized supply-to-demand ratio Φ is

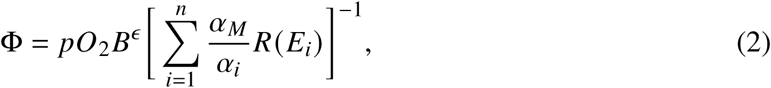

where the *α*_*i*_ represent the supply rate coefficients at reference temperature and 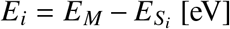 [eV] denote the differences between the sensitivities of metabolic demand and the supply processes. The dependence of supply and demand on body mass *B* is reflected in the allometric exponent *ϵ* as in the original index (6).

The condition for the existence of a bowl-shaped *P*_*crit*_ curve, i.e. supply steps having temperature sensitivities both less than and greater than that of metabolic demand, thus reads 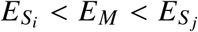 for any two supply steps *i* and *j*. Eqn. (2) can include any number of supply processes. However, we find that *P*_*crit*_ curves generated by the full model (*n* = 3) can still be appropriately represented by curves assuming only 2 steps. Adding more exponential curves does not change the qualitative range of possible *P*_*crit*_ curves beyond those of concave-up bowl shapes (Fig. S4A and C). The generalized Metabolic Index in Eqn. (2) can also reproduce *P*_*crit*_ curves that include the Hx/HxO system (Fig. S4B, D and E), because the effects of the chemical blood component on the *P*_*crit*_ curve are qualitatively the same as those of the biophysical parameters (Fig. 2C and D). In such cases, however, the parameters can no longer be associated with single steps in the supply chain, but instead capture the combined properties of the processes that limit the O_2_ supply towards the cold and warm ends of the temperature range, respectively.

### Connecting Physiology to Biogeography

The metabolic index framework establishes a direct link between physiological traits and biogeographical distributions, as the range boundaries of a diverse set of species align more strongly with a specific value of the index than with either temperature or pO_2_ alone (7). The generalized formulation has the potential to further improve this description of species habitats, especially at the cold edges of a species distribution.

To examine whether the thermal optima in physiological hypoxia tolerance are reflected in a species’ biogeography, we investigate state-space habitats of biogeographic occurrence data from the Ocean Biodiversity Information System (35, Materials and Methods). For the species presented in Fig. 1, the environmental habitat conditions are poorly represented in large-scale datasets (Fig. S6). However, for two additional species with physiological traits suggesting a thermal optimum, the starry flounder *Platichthys stellatus* and the shrimp *Oplophorus spinosus*, adequate occurrence and environmental data are available. In *Platichthys stellatus*, estimates from published experimental results yield a temperature sensitivity *E*_*M*_ = 0.68 eV for metabolism and *E*_*Vent*_ = 0.9 eV (36) for the ventilation rate, indicating a bowl-shaped O_2_ limitation. For *Oplophorus spinosus*, critical O_2_ pressures have been measured and display a minimum at intermediate temperatures (37), such that Eqn. (2) can be fit directly.

In both species, the environmental conditions in occupied habitats reveal a clear minimum in inhabited pO_2_ at intermediate temperatures, consistent with the physiological predictions (Fig. 5). In contrast, the minimum inhabited temperatures of each species are inconsistent with a model based on a lower threshold value of temperature that is independent of O_2_. Instead, minimum temperatures decrease to lower values as oxygen levels increase. Similar patterns are also observed in other species for which laboratory experiments indicate thermal optima and for which sufficient occurrence data are available (SI, Fig. S7, 16).

**Fig. 5.**
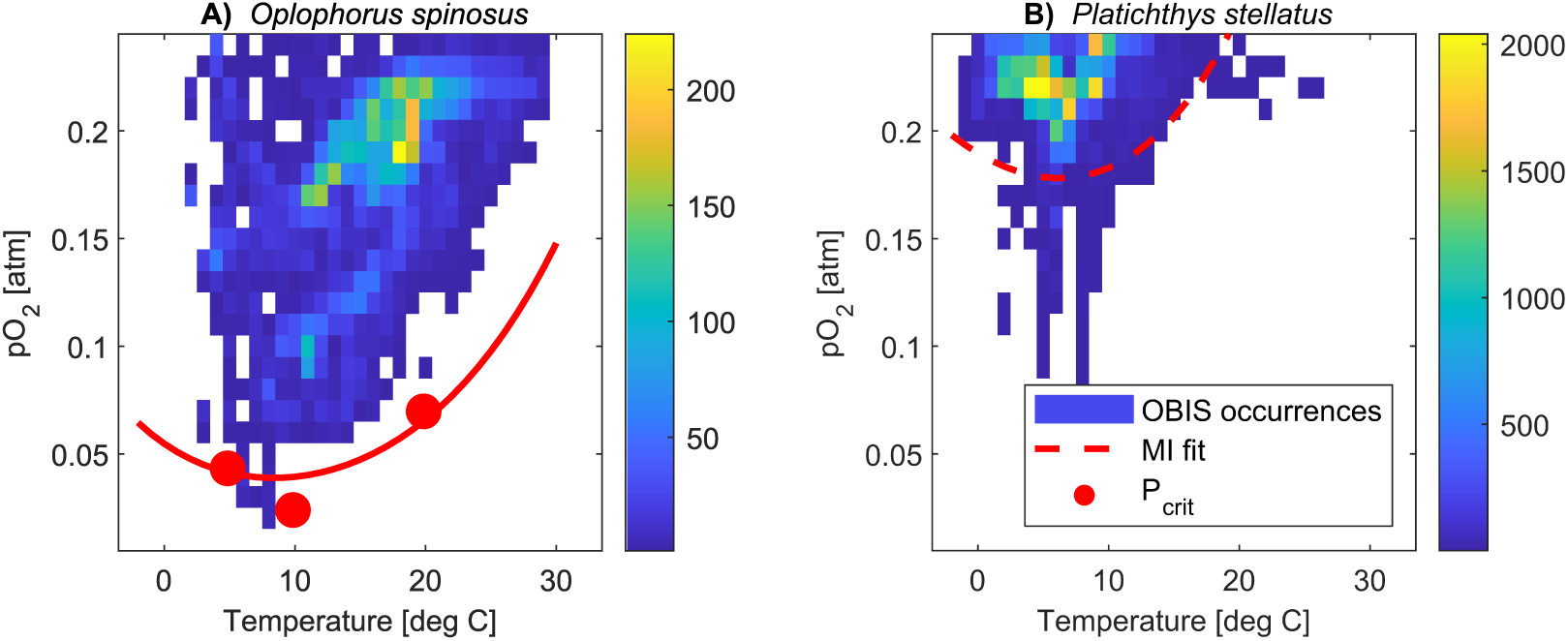
Temperature and pO_2_ state-space habitats using global occurrence data from the Ocean Biodiversity Information System reveal a thermal optimum at intermediate temperatures in agreement with physiological evidence. **A)** In the midwater shrimp *Oplophorus spinosus*, measured *P*_*crit*_ values (red dots) as well as the curve fit based on the Metabolic Index (solid red line) align with the lowest inhabited pO_2_ across the temperature range. **B)** In the flounder *Platichthys stellatus*, the *P*_*crit*_ curve (dashed red) predicted from the Metabolic Index framework and physiological rates also exhibits a thermal optimum consistent with the occurrence data.

In all these cases, the generalized Metabolic Index reveals how the reversal in hypoxia tolerance at low temperatures results from physiological traits, and how this bidirectionality is reflected in biogeographic ranges. In particular, it suggests O_2_ limitation is the mechanism that restricts habitat towards the cold edges of species distributions.

## Discussion

The dynamic model of temperature-dependent hypoxia reveals that a series of biophysical O_2_ supply steps can give rise to thermal optima in hypoxia tolerance as observed in new respirometry data. This occurs when the supply chain includes at least two processes such that one accelerates with temperature more slowly than metabolic demand, and another accelerates more rapidly. In this case, the process with a lower temperature sensitivity drives an increase in *P*_*crit*_ under warm conditions, while the more sensitive process leads to a reversal with higher *P*_*crit*_ in cold waters. A generalized Metabolic Index adequately captures these complex patterns in a single metric based on mechanistic principles.

Our analysis of available physiological evidence suggests that such bidirectional effects of temperature on hypoxia tolerance may not be uncommon in aquatic animals across taxonomic groups. Estimates of the temperature sensitivity of ventilation and circulation rates in aquatic ectotherms fall above diffusive gas exchange and below metabolism on average, but imply the existence of thermal optima in a significant fraction of species. However, these results rely on limited physiological data. In particular, there are only a few teleost and crustacean species for which all required physiological estimates are available. Thus, sampling the involved traits across a broader range of the taxonomic, morphological and ecological diversity is a key step towards further advancing and testing this framework and its implications.

In contrast to the sparsity of detailed physiological measurements, global occurrence data is available for a much larger number and diversity of marine species (e.g OBIS). The generalized index offers further improvements in the analysis of these data compared to its original formulation, especially along the cold edges of species habitats by including a meaningful representation of O_2_ limitation at low temperatures. In case studies presented here, thermal optima in physiological hypoxia tolerance are also reflected in species’ biogeographic state space. Leveraging this approach in a future database-wide analysis of occurrence data will contribute to a fuller picture of how temperature and oxygen shape the biogeography and ecology of marine species.

Oxygen limitation of aerobic metabolism at low temperature has broad implications for marine ecosystems and their response to climate change. Marine species richness is generally observed to decline towards the poles, and is often cited as being driven by gradients in ocean temperature, with cooler waters taken to inhibit diversity (38). Our results indicate that long-term aerobic energy constraints on viable habitat in cold water could be a physiological cause of this poleward diversity loss. At the same time, warming at species’ poleward range limits would relieve such aerobic constraints, allowing species to disperse towards, and establish in, higher latitudes. This mechanism could thus potentially explain widespread poleward migrations of marine species seen in response to recent anthropogenic warming (39, 40). On longer timescales, O_2_ limitation at species’ cold edge habitat limits provides a novel mechanism for driving habitat loss during periods of global cooling, and may underlie previous extinctions during such phases (41, 42).

## Materials and Methods

### Laboratory Measurements

Critical O_2_ levels were measured following standard closed system respirometry protocols (17, 43) for individuals of *T. tubifex* (*n* = 132), *N. vectensis* (*n* = 107) and *L. pictus* (*n* = 40). For the social squid *D. opalescens*, we measured critical O_2_ levels for 14 groups of 15 to 30 (median 20) animals following published closed system respirometry protocols for this species (44). *P*_*crit*_ was determined by breakpoint analysis of the O_2_ draw down curve (45). Full protocols are provided in the SI.

### Dynamic Model

The pools and fluxes of O_2_ in a generic water-breather are described by a nonlinear system of 8 ordinary differential equations. For each set of model parameters, simulations are performed across the temperature range from 0 °C to 30 °C until *P*_*crit*_ can be determined by breakpoint analysis from the rate of O_2_ draw down. All simulations were carried out in the Python language using the solve_ivp function in Scipy (46) for numerical integration. The full model description is provided in the SI.

### Ventilation and Circulation Data

We compiled data on ventilation rates (*n* = 8), ventilation frequency (*n* = 18), ventilation stroke volumes (*n* = 6), circulation rates (*n* = 4), heart rates (*n* = 20) and heart stroke volumes (*n* = 2) of aquatic water-breathers measured at two or more temperatures at atmospheric O_2_ levels. Estimates of the sensitivity parameters *E*_*V*_, *E*_*C*_ were obtained through least square fits of Arrhenius functions to the experimental data using the curve_fit function and density estimates were obtained using the gaussian_kde function in Scipy. A detailed description of the compiled data is provided in the SI and all estimates are availabe in Dataset S1.

### State-space Habitats

State-space habitats were obtained by pairing species location data downloaded from the Ocean Biodiversity Information System (35) in September 2019 with monthly temperature and O_2_ conditions from the World Ocean Atlas (47, 48) according to the procedure described in (7). All available state-space habitats are shown in Fig. S6 and Fig. S7. Additional information is provided in the SI.

## Acknowledgments

The study was made possible by grants to C.D. from the National Oceanic and Atmospheric Administration (NOAA NA18NOS4780167), the California SeaGrant and Ocean Protection Council, and the National Science Foundation (NSF OCE-1737282). E.A.S. is funded by NSF EAR-1922966 and an Environmental Ventures Project grant from the Stanford Woods Institute. We thank Leanne Elder, Pincelli Hull, and Murray Duncan for helpful discussion, and Andy Marquez for laboratory assistance.

## Data and Code Availability

Sensitivity estimates for ventilation and circulation rates obtained from published results are available in Dataset S1. Biogeographic and environmental data are publicly available. Full physiological measurements are shown in Fig. S1. and are available from the corresponding author upon reasonable request. Python code will be made available upon publication.

## Notes

### Competing Interest Statement

The authors have declared no competing interest.

