## Supplementary Information for "Physiological causes and biogeographic consequences of thermal optima in the hypoxia tolerance of marine ectotherms"

**This Supplementary Information includes:**

Supplementary Text

Figures S1 to S7

Table S1

Legend for Dataset S1

SI References

**Other supplementary materials for this manuscript include the following:**

Dataset S1

### Supplementary Information Text

#### A. Respiration physiology experiments

Full $P_{crit}$ data for all species are shown in Fig. S1.

*Nematostella vectensis* were lab reared and collected in 2017 at Stanford Hopkins Marine Station in Pacific Grove, California. *Lytechinus pictus* were purchased from the South Coast Bio-Marine Biological Supply in San Pedro, California, and analyzed in 2019. Collection of each species was seasonally timed so that the experimental temperature range was as close as possible to the current environmental temperature range. Upon arrival to the lab, animals were transferred into a synthetic seawater (33 practical salinity units, PSU) aquarium. Each animal was isolated, starved and acclimated to experimental temperature (± 0.3 °C) for a minimum of 12 hours (up to 24 hours ), then individually placed into a darkened respiration chamber with an oxygen optode attached to a Pyroscience FireSting 4-channel optical oxygen meter. Oxygen optodes were calibrated at each experimental temperature using a saturated sodium sulfite ($Na_2SO_3$) solution (30 g $L^{-1}$) for 0% oxygen saturation, while a 100% oxygen saturation was achieved using seawater fully aerated with an air stone. A separate reservoir of synthetic seawater was used to fill experimental chambers prior to runs to ensure metabolites and microbes were absent prior to each run. Glassware was autoclaved between runs, and at temperatures above 25 °C synthetic seawater was UV-treated and an antibiotic Streptomycin sulfate salt ($C_{21}H_{39}N_7O_{12}\cdot$ 1.5z$H_2SO_4$) solution (50 $\mu$mol $L^{-1}$) was also added to further inhibit bacterial respiration. Bacterial respiration was negligible and was tracked using a blank respiration chamber that was filled with identically treated synthetic seawater. At the end of experiments specimens were blotted dry and weighed to obtain wet mass. Standard closed system respirometry protocols were used to measure $P_{crit}$. The oxygen content of each chamber was continuously recorded at one second intervals using Pyro Oxygen Logger software, allowing $P_{crit}$ to be determined with break-point analyses on the oxygen draw down curve (1). This is possible because $P_{crit}$ coincides with the onset of anaerobiosis, representing the switch from oxyregulatory aerobic respiration independent of $pO_2$, to oxyconformity, whereby the animal slows respiration in response to decreasing $pO_2$. This is reflected in a breakpoint between slopes representing oxygen consumption (standard metabolic rate), and drastically reduced oxyconformation oxygen consumption as anaerobic metabolism makes up an increasingly larger fraction of ATP turnover. To determine the effect of temperature on the metabolic rate and hypoxia tolerance, experiments were repeated across five temperatures which span natural environmental range. A minimum of 10 individuals were measured for each temperature to account for natural variation, with 107 *Nematostella vectensis* and 40 *Lytechinus pictus* individuals measured in total across all temperatures. Resting $MO_2$ at different temperatures was determined from the slope of the oxygen drawdown curve in each experimental chamber prior to $P_{crit}$ and incorporating the adjusted volume of water, mass of the organism, and time elapsed using the following equation (2):

$$MO_2 = \frac{\frac{[O_2]_2-[O_2]_1}{t_2-t_1} \cdot 3600 \cdot V}{B}$$

where

$MO2$ = mass-normalized metabolic rate ($\mu$mol $L^{-1}$ $g^{-1}$ $h^{-1}$)
$[O_2]$ = oxygen concentration ($\mu$mol $L^{-1}$)
$t$ = time (s)
$V$ = water volume (L)
$B$ = body mass (g)

To determine standard metabolic rate at each temperature, recorded MO2 measurements were resampled 10,000 times to produce a frequency distribution. The mean of the lowest 10% of points was used while all MO2 measurements above this were assumed to be associated with spontaneous activity (3).

We used closed-system respirometry to measure the temperature dependence of $P_{crit}$ in groups of *D. opalescens*. Experiments controlled for other factors known to influence resting and routine levels of metabolic demand of marine ectotherms: body size, digestion, and oxygen availability (4). Adult *D. opalescens* were captured in nearshore waters of Monterey Bay, CA, USA from April-May 2018 using small barbless jigs and then transported in aerated seawater to an indoor aquarium facility at Hopkins Marine Station, Pacific Grove, CA. Only undamaged squid caught by the sucker cups on the arms were retained for this study. No injuries were observed in any animals used in experiments (n = 309; 187 male, 111 female). Squid mass ranged from 17.7-89.1 g, with an average of 41.4 g. Squid were housed together in groups in a 3200 L circular tank with filtered flow-through seawater (20 L $min^{-1}$) at ambient temperature (12-14 °C) for 24-48 h during which they received one meal of live minnows (*Pimephales promelas*) per 24 h period. Group respiration was accomplished using a custom-built respirometer. The system is composed of two 600 L circular tanks arranged side-by-side a buffer tank for temperature control and an insulated test tank for metabolic measurements (484 L test section when sealed with a transparent lid). The test tank was visually shielded with opaque plastic sheeting suspended over a fence encircling the circumference, but squid could be observed through a small opening. All experiments occurred at the same time of day and under the same lighting conditions. After introduction to the test tank, groups of squid were allowed to acclimate to experimental conditions for 12 h with flow-through seawater (range = 12.5-13.2 °C, mean = 13 °C). This resulted in at least 24 h without food before measurement began. We used a HQ40D multimeter and LDO101 oxygen sensor (Hach Co., USA) to record oxygen concentration, oxygen saturation, and temperature at 0.1 Hz in the test tank. During the temperature ramp, an immersion heater and immersion chiller (Process Technology, USA) connected to the buffer tank respectively increased or decreased the water temperature at 1 °C $h^{-1}$ as measured in the test tank. Once the temperature ramp was complete, measurement began and continued until squid started showing obvious signs of fatigue, or until oxygen concentrations reached 0.5 $mgL^{-1}$ (measurement duration range = 6.4-23.9 h, median = 11.9 h). Measurements were conducted at a suite of temperatures within the ecologically relevant range that *D. opalescens* typically encounters in the CCS. Three groups were measured at each of the following treatments: 7.5, 10, 13, 16, and 19 °C. Due to a failure in power supply to the multimeter, data from one experiment at 16 °C were not recorded and thus there was a total of n = 14 groups. The average temperature during measurement was never more than 0.4 °C from the target temperature. The routine activity *D. opalescens* displayed during measurements, where squid used a mixture of fin undulation and jet propulsion to slowly move head- and tail-first and to hold station, sufficiently mixed water within the test tank. Using functions in the respR package in R (5), we applied the segmented regression approach (1) to determine the breakpoint of oxygen consumption curves and calculate $P_{crit}$, the oxygen partial pressure below which a stable rate of oxygen uptake at routine levels was not maintained (6). Oxygen consumption rates were consistent from experiment start until $P_{crit}$ was reached. Control experiments containing no squid were conducted to quantify microbial background respiration. No background respiration was detected over experimentally relevant durations; we therefore did not apply any correction to group rates for background respiration.

#### B. Dynamic model formulation

The time-dependent transfer of $O_2$ from the environment to the metabolizing tissues is simulated using a system of 8 ODEs representing the concentrations of dissolved $O_2$ $(X)$ as well as the free $(F)$ and bound $(B)$ forms of $O_2$ transporting proteins in the four model compartments as shown in Fig. S2.

The full ODE system reads:

$$\text{Dissolved O}_2\ X\ :\begin{cases} V_W \frac{d}{dt} X_W &= -\hat{\alpha}_V (X_W - X_B) \\ V_B \frac{d}{dt} X_B &= \hat{\alpha}_V (x_W - X_B) - \hat{\alpha}_D \frac{(X_B - X_G)}{\kappa_H} \\ V_G \frac{d}{dt} X_G &= \hat{\alpha}_D \frac{(X_B - X_G)}{\kappa_H} - \hat{\alpha}_C (X_G - X_I) + k_b B_G - k_f X_G^n F_G \\ V_I \frac{d}{dt} X_I &= \hat{\alpha}_C (X_G - X_I) + k_b B_I - k_f X_I^n F_I - \hat{\alpha}_M \end{cases} \tag{1-4}$$

$$\text{Free transport protein } F\ :\begin{cases} V_G \frac{d}{dt} F_G &= -\hat{\alpha}_C (F_G - F_I) + k_b B_G - k_f X_G^n F_G \\ V_I \frac{d}{dt} F_I &= \hat{\alpha}_C (F_G - F_I) + k_b B_I - k_f X_I^n F_I \end{cases} \tag{5-6}$$

$$\text{O}_2 - \text{bound transport protein } B\ :\begin{cases} V_G \frac{d}{dt} B_G &= -\hat{\alpha}_C (B_G - B_I) - k_b B_G + k_f X_G^n F_G \\ V_I \frac{d}{dt} B_I &= \hat{\alpha}_C (B_G - B_I) - k_b B_I + k_f X_I^n F_I \end{cases} \tag{7-8}$$

Here $X_W, X_B, X_G, X_I$ denote the concentrations of dissolved $O_2$ $[\frac{\mu mol}{l}]$ in the external water, external boundary layer, internal gill and internal tissue compartments with corresponding volumes $V_W, V_B, V_G, V_I$ $[l]$, while $F_G, F_I$ and $B_G, B_I$ denote the concentrations of free and $O_2$-bound transport proteins $[\frac{\mu mol}{l}]$ in the gill and tissue compartments, respectively. The transport of $O_2$ depends on temperature $T[K]$ and is realized through ventilation $\hat{\alpha}_V(T)[\frac{l}{min}]$ between the environmental water and the boundary layer, through diffusive gas exchange $\hat{\alpha}_D(T)[\frac{\mu mol}{min\ kpa}]$ between the boundary layer and the gill volume and through circulation $\hat{\alpha}_C(T)[\frac{l}{min}]$ between the gill volume and the tissues. For diffusive gas exchange, the concentration difference between the compartments is converted to a partial pressure difference with the temperature-dependent solubility $\kappa_H(T)$ $[\frac{\mu mol}{l\ kpa}]$ of $O_2$ in seawater, calculated using coefficients from (7) for (8) for a salinity of 35 PSU, assuming that the relative solubility of $O_2$ in blood does not change with temperature for the purposes of this model (while $O_2$ solubility is lower in body fluid than in seawater, e.g. (9), any temperature-independent effect only changes the rate constant of diffusion). Finally, $O_2$ is consumed in the tissue compartment through metabolism $\hat{\alpha}_M$ $[\frac{\mu mol}{min}]$.

These supply and demand rates $\hat{\alpha}_V, \hat{\alpha}_D, \hat{\alpha}_C, \hat{\alpha}_M$ are defined as

$$\begin{aligned}\hat{\alpha}_V(T) =& \alpha_V \cdot R(T, E_V) \\ \hat{\alpha}_D(T) =& \alpha_D \cdot R(T, E_D) \\ \hat{\alpha}_C(T) =& \alpha_C \cdot R(T, E_C) \\ \hat{\alpha}_M(T) =& \alpha_M \cdot R(T, E_M)\end{aligned} \tag{9-12}$$

with constants $\alpha_x$ and dimensionless temperature dependent factors taking the form of Arrhenius functions

$$R(T, E_x) = \exp\left(\frac{-E_x}{k_B}(\frac{1}{T} - \frac{1}{T_{ref}})\right) \tag{13}$$

with temperature sensitivities $E_x$ $[eV]$. Throughout the paper, we use the reference temperature $T_{ref} = 288.15$ $[°K] \equiv 15$ $[°C]$ and $k_B$ denotes the Boltzmann constant. The temperature-independent $\alpha_x$ represent the supply and demand fluxes at the reference temperature determined by biological traits like resting metabolic rate, gill surface area, heart rate etc. These rate coefficients and temperature sensitivities are varied throughout the investigation to explore their impacts on hypoxia tolerance. For diffusion, we estimate $E_D$ by fitting an Arrhenius function to relative diffusivity ($\kappa$) values

$$\frac{\kappa(T)}{\kappa(T_{ref})} = \frac{T}{T_{ref}} \frac{\mu(T_{ref})}{\mu(T)} \tag{14}$$

with the dynamic viscosity of seawater $\mu(T)$ calculated according to (10) for a salinity of 35 PSU. After accounting for temperature-dependent solubility $\kappa_H$, this yields a value of $E_D \approx 0.06$ eV.

Typical values of the biophysical parameters and values used for the compartment volumes are provided in Table S1.

The model also contains a blood component with free ($F$) and bound ($B$) forms of transport protein circulating between the gill and the tissue compartments according to $\hat{\alpha}_C(T)$. These bind or release $O_2$ with forward and backward reaction rates $k_f$ $[\frac{l^3}{\mu mol^2\ min}]$, $k_b$ $[\frac{l}{min}]$, respectively, with a dissociation curve following Hill's equation where the $O_2$ pressure producing half-saturation, denoted $P_{50}$, is temperature-dependent (11). The fraction $Y$ of protein bound to $O_2$ is

$$Y = \frac{[B]}{[B] + [F]} = \frac{pO_2^{n_H}}{P_{50}^{n_H} + pO_2^{n_H}} \tag{15}$$

where

$$P_{50}(T) = P_{50}^{T_{ref}} \cdot \exp\left(\frac{\Delta H}{R_{Gas}}(\frac{1}{T} - \frac{1}{T_{ref}})\right) \tag{16}$$

with the half-saturation pressure at the reference temperature $P_{50}^{T_{ref}}$, the Hill coefficient $n_H$ and the enthalpy of the binding reaction $\Delta H$, and the gas constant $R_{Gas}$. The $O_2$ concentration $K_{50}$ producing half saturation ('microscopic dissociation constant') can be converted from the pressure $K_{50} = \kappa_H P_{50}$. Since the ratio of the reaction rates $k_f, k_b$ determining the equilibrium is more relevant than the absolute values of the rates for our purposes, we fix the forward rate to be much faster than diffusive gas exchange and compute $k_b$ as $k_b = k_f K_{50}^{n_H}$ from the equilibrium constant $K_{eq} = \frac{k_b}{k_f} = K_{50}^{n_H}$).
The values used for these chemical parameters are shown in Table S1.

At each temperature, we integrate the ODE system using the *solve_ivp* solver in Python (12). We start each simulation with a concentration of dissolved $O_2$ typical of surface seawater of

300 $\frac{\mu mol}{l}$ in all compartments and with corresponding equilibrium concentrations of the transport protein. After an initial spin-up time of 50 min with constant $X_W(t)$ to ensure that the saturated $O_2$ concentration can support the model metabolic rate, $X_W(t)$ starts to evolve according to SI Eqn. (1). To determine $P_{crit}$, we use the breakpoint analysis (13) by applying a segmented linear regression following (1) to the rate of change of $X_W(t)$. We use the time $t_{crit}$ at which the $O_2$ concentration in the tissue first drops below a critical limit near zero ($X_I(t_{crit}) \leq 0.05 \frac{\mu mol}{l}$) as an initial guess for the single breakpoint and restrict the regression to the interval $[50, t_{crit} + \frac{t_{crit}}{5}]$. If $X_I$ never drops below this threshold during the default simulation time of 1000 min, the time is doubled and the process is repeated until a $P_{crit}$ value is obtained.

To illustrate the diminishing effects of supply processes that accelerate faster than metabolism under warm condition, we formulate a simple model variant with a modified ventilation rate with a slowed acceleration above 20 °C:

$$\hat{\alpha}_V = \begin{cases} \alpha_V R(T, E_V) & T < 288.15\ K \\ \frac{2\alpha_V}{1+\exp(-\lambda(T-T_{ref})} & T \geq 288.15\ K \end{cases} \tag{17}$$

where $\lambda = \frac{2E_V}{k_B T_{ref}^2}$ is chosen to match the value and slope of the Arrhenius ventilation at $T_{ref}$. The modified rate as well as the resulting change in $P_{crit}$ is shown in Fig. S3.

#### C. Metabolic Index derivation

The generalized Metabolic Index (Eqn. (2) in main text) can be derived from the dynamical model without the blood proteins $F, B$ in SI Eqn. (1) to (4). To determine $P_{crit}$ analytically, we consider the case of a very large environmental volume $V_W \to \infty$, which is equivalent to a setting with a constant environmental $O_2$ level $X_W(t) = X_W$ that is not depleted over time. Intuitively, this represents the fact that respiration of a small model organism does not significantly change the ambient $O_2$ level in the ocean or a large tank. Under this assumption, we can find the constant concentration $X_W^{crit}$ that results in a steady state $O_2$ level of zero in the tissue, $X_I^{ss} = 0$, which corresponds to the criterion for reaching $P_{crit}$ in our model. From SI Eqn (4), we have

$$\frac{d}{dt}X_I = \hat{\alpha}_C(X_G^{ss} - 0) - \hat{\alpha}_M = 0$$
$$\Rightarrow X_G^{ss} = \frac{\hat{\alpha}_M}{\hat{\alpha}_C}.$$

Substituting this in SI Eqn 3 yields

$$\frac{d}{dt}X_G = \hat{\alpha}_D \frac{(X_B^{ss} - \frac{\hat{\alpha}_M}{\hat{\alpha}_C})}{\kappa_H} - \hat{\alpha}_M = 0$$
$$\Rightarrow X_B^{ss} = \frac{\hat{\alpha}_M \kappa_H}{\hat{\alpha}_D} + \frac{\hat{\alpha}_M}{\hat{\alpha}_C}.$$

Finally, inserting $X_B^{ss}$ and $X_G^{ss}$ in SI Eqn. (2) yields

$$\frac{d}{dt}X_B = \hat{\alpha}_V(X_W^{crit} - \frac{\hat{\alpha}_M \kappa_H}{\hat{\alpha}_D} - \frac{\hat{\alpha}_M}{\hat{\alpha}_C}) - \hat{\alpha}_M = 0$$
$$\Rightarrow X_W^{crit} = \frac{\hat{\alpha}_M}{\hat{\alpha}_V} + \frac{\hat{\alpha}_M \kappa_H}{\hat{\alpha}_D} + \frac{\hat{\alpha}_M}{\hat{\alpha}_C}.$$

Converting to partial pressure results in the expression for $P_{crit}$

$$P_{crit} = \frac{\hat{\alpha}_M}{\hat{\alpha}_V \kappa_H} + \frac{\hat{\alpha}_M}{\hat{\alpha}_D} + \frac{\hat{\alpha}_M}{\hat{\alpha}_C \kappa_H} \tag{18}$$

This corresponds to the MI

$$\Phi = pO_2 B^{\epsilon} \left( \frac{\hat{\alpha}_M}{\hat{\alpha}_V \kappa_H} + \frac{\hat{\alpha}_M}{\hat{\alpha}_D} + \frac{\hat{\alpha}_M}{\hat{\alpha}_C \kappa_H} \right)^{-1} \tag{19}$$

which matches Eqn. (2) in the main text after substituting SI Eqns. (9) to (12), including the body mass dependence $B^{\epsilon}$ of the original index. On the other hand, the same result is obtained when considering the ratio of $O_2$ supply to demand directly, since the total (potential) supply $S_{tot}$ of the supply chain is

$$S_{tot} = pO_2 \cdot \hat{\alpha}_{tot}$$

with the total conductance of the linear chain

$$\hat{\alpha}_{tot} = \left( \sum_{i=1}^{n} \frac{1}{\hat{\alpha}_i} \right)^{-1}$$

Now the ratio $\frac{S_{tot}}{\hat{\alpha}_M}$ once again yields SI Eqn. (19).

While the full system including the transport protein concentrations $F, B$ does not yield a simple expression for $P_{crit}$ and $\Phi$, simulated $P_{crit}$ curves can be adequately fit based on Eqn. 2 in the main text with only $n = 2$ supply steps, as shown in Fig. S4 for a range of biophysical and chemical parameters. In particular, the figure illustrates the fact that the presence of $O_2$ transporting proteins essentially enhances the supply capacity of circulation in the model, with changes in the total concentration of protein achieving a similar effect to changes in the parameter $\alpha_C$ (Fig. S4 A and B) and changes in the enthalpy of the binding reaction corresponding to changes in the sensitivity $E_C$ (Fig. S4 C, D).

### D. Ventilation and circulation data

We compiled data on the thermal sensitivity of ventilation rates (n=8), ventilation frequencies (n=18), ventilation stroke volumes (n=6), circulation rates (n=4), heart rates (n=20) and heart stroke volumes (n=2) in aquatic water breathers at atmospheric $O_2$ levels measured at 2 or more temperatures from the literature. For each of the 58 datasets from 35 species (25 marine, 10 freshwater), the temperature dependence of the measured quantity was estimated by fitting an Arrhenius equation $k \cdot R(T, E)$ from SI Eqn. (13) to the data normalized to the lowest experimental temperature and estimating the sensitivity parameter $E$. Least-Square Fits were performed using the *scipy.optimize.curve_fit* function. The results are provided in Dataset S1.

The temperature sensitivities of ventilation (32 estimates) and circulation (26 estimates) were not significantly different in terms of means according to a F-test ($p = 0.1608$), which was performed after testing for equal variances (Levene-test, $p = 0.8103$) and checking normality (Shapiro-Wilk-test, $p = 0.5071$ and $p = 0.5816$, respectively). Therefore, the ventilation and circulation estimates were combined for further analysis.

We estimated the expected frequency of the supply temperature sensitivities $E_V, E_C$ of volumetric flow rates exceeding the sensitivity of metabolism $E_M$. To that end, we estimated the trait frequency distributions of these sensitivities through kernel density estimation using the gaussian_kde function in Scipy and sampled from the resulting densities ($n = 100000$). To account for the effect of decreasing solubility $\kappa_H$ with temperature as it appears in SI Eqn. (18), the density of ventilation and circulation rates was shifted by the temperature sensitivity of solubility ($\approx 0.144$ eV from 8). This procedure yields an expected frequency for $E_V, E_C > E_M$ of 23 %, which corresponds to the expected fraction of species with thermal optima in hypoxia tolerance if the traits are considered independent.

On the other hand, in the 17 species for which both estimates are available, we find that $E_V, E_C > E_M$ in 7 cases (frequency $\approx 41$ %) after accounting for solubility effects, indicating that the traits determining the thermal sensitivity of $O_2$ supply and demand are unsurprisingly not independent in real organisms. The temperature sensitivities in these 17 species are comparable to estimates obtained from all species (demand 0.58 ± 0.15 eV, supply 0.43 eV ± 0.21 eV, Fig. S5A).

To evaluate whether $E_V, E_C$ are themselves temperature dependent, we estimated the parameter separately above and below the median temperature in the 40 datasets from 25 species with 4 or more distinct experimental temperatures. Estimates were again obtained by fitting Arrhenius functions $k \cdot R(T, E)$ to the data. For the 10 estimates of volumetric flow rates, the cold side mean of 0.69 eV far exceeds the warm side mean of 0.07 eV (paired sample t-test, $p = 0.009$). The difference remains significant when frequencies and stroke volumes are also considered (cold side mean 0.43 eV, warm side mean of 0.15 eV, paired sample t-test, $p = 0.0085$). This is illustrated in Fig. S5B. If the analysis is restricted to the 24 datasets from 16 species with temperature ranges of at least 5 °C both above and below the median experimental temperature, the difference between the cold side mean (0.48 eV) and the warm side mean (0.22 eV) remains similar. The average experimental temperature range is 17.95 °C (10.95 °C cold side, 7.51 °C warm side).

#### E. State-space projections

The curves in Fig. 5 were obtained from Eqn. (2) with $n = 2$ supply steps while accounting for any physiological evidence available for the respective species.

For *Oplophorus spinosus*, Eqn. (2) was fit directly to the experimental $P_{crit}$ data (14) while constraining the temperature sensitivities to $0 < E_1 < 0.8$ [eV] and $-0.7 < E_2 < 0$ [eV] respectively, based on the estimate $E_M \approx 0.7308$ eV estimated in (15) from the data of (14).

For *Platichthys stellatus*, no estimates of $P_{crit}$ are available, so the curve was tuned to $E_1 = 0.65$ eV and $E_2 = -0.25$ eV based on the available estimates of $E_{Vent} \approx 0.9$ eV and $E_M \approx 0.68$ eV, as well as the theoretical value of $E_D \approx 0.06$ eV for diffusion.

In addition to the state-space habitats shown in Fig. 5, we also provide the habitats for the 4 species with experimental $P_{crit}$ data (Fig. 1) in Fig. S6. There are only very few occurrences available in OBIS for three of the species (*L. pictus, N. vectensis, T. tubifex*), two of which are intertidal (*N. vectensis*) or inhabit freshwater and brackish habitat (*T. tubifex*). In these cases, the use of monthly mean values for temperature and $O_2$ conditions from the World Ocean Atlas likely does not reflect the range of environmental conditions these species experience in their natural habitats, such as strong variations in $O_2$ frequently experienced by intertidal animals (16), and limits the interpretation of the state-space projections. While there is a larger number of occurrences for *D. opalescens* (Fig. S6A), the gradients in temperature and $O_2$ these squid experience during offshore migrations and nearshore spawning activity (17) are likely also not well represented in monthly climatologies.

Finally, we provide the state-space habitats for the 8 marine species for which both estimates of the supply and demand temperature sensitivities are available (Fig. S7, Dataset S1). The state-space habitats reveal bowl-shaped patterns in the two additional marine species for which physiological evidence suggests the presence of thermal optima (*C. crangon, S. solea*). Such bowl-shaped patterns are also present in the state-space habitats of some of the remaining species, for which the available estimates only explain the warm edge (e.g. *G. morhua*, Fig. S7E). However, there is no species with all required sensitivity estimates for metabolic demand, ventilation rate and circulation rate to test the emergence of state-space habitats without thermal optima.

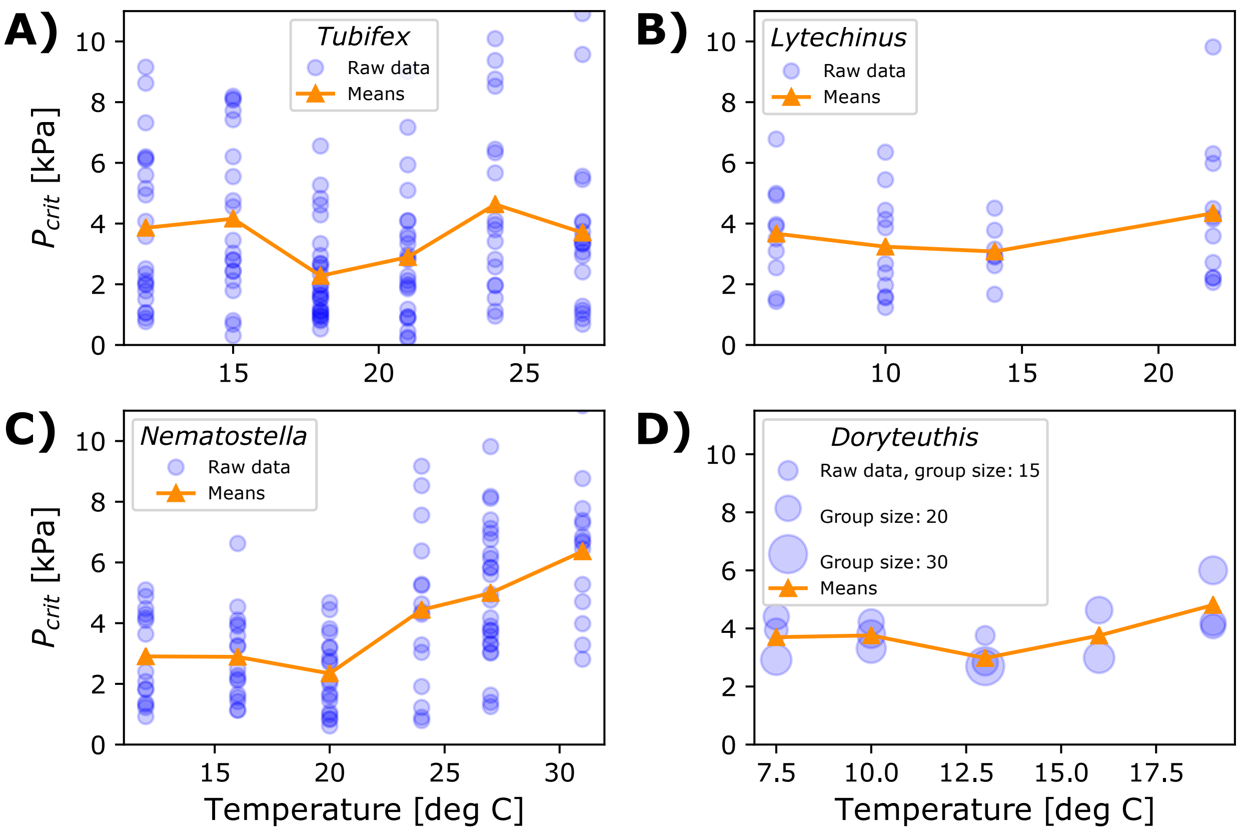


273 Fig. S1. Raw $P_{crit}$ measurements (blue) and mean values as in Fig. 1 (orange) for **A)** the oligochaete worm *Tubifex tubifex*, **B)** the sea urchin *Lytechinus pictus*, **C)** the anemone *Nematostella vectensis* and **D)** the market squid *Doryteuthis opalescens*. Experimental protocols and the procedure for determining $P_{crit}$ values are provided in SI Text A.

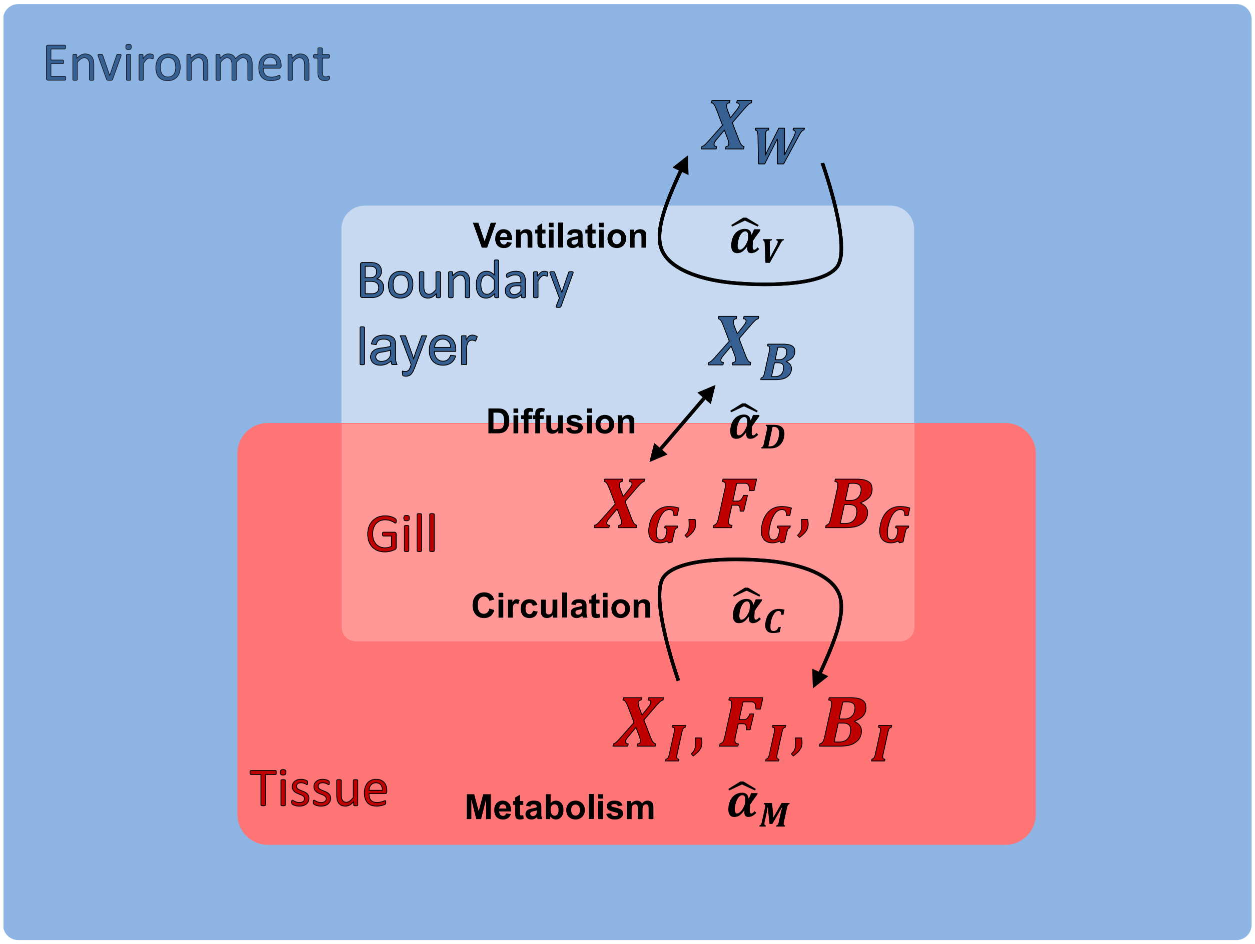


Fig. S2. Illustration of state variables for dissolved $O_2$ ($X$), free ($F$) and bound ($B$) $O_2$-transport protein as well as temperature-dependent $O_2$ supply and demand processes ($\hat{\alpha}$) used in the dynamical ODE model.

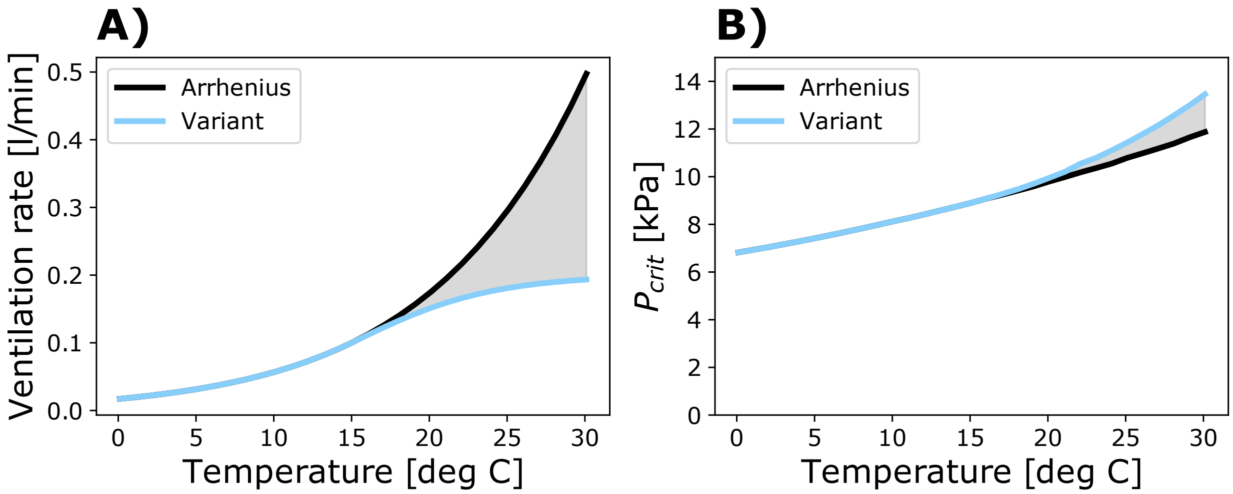


Fig. S3. Variant of the dynamical model featuring a ventilation rate with slower acceleration at high temperatures. **A)** Above 20 °C, the modified ventilation rate following SI Eqn. (17) accelerates significantly slower than the default rate governed by the exponential (Arrhenius) relationship in SI Eqn. (9). **B)** The resulting change in $P_{crit}$ at high temperatures is minimal because ventilation is characterized by a larger temperature sensitivity than metabolism ($E_V = 0.8eV, E_M = 0.5eV$).

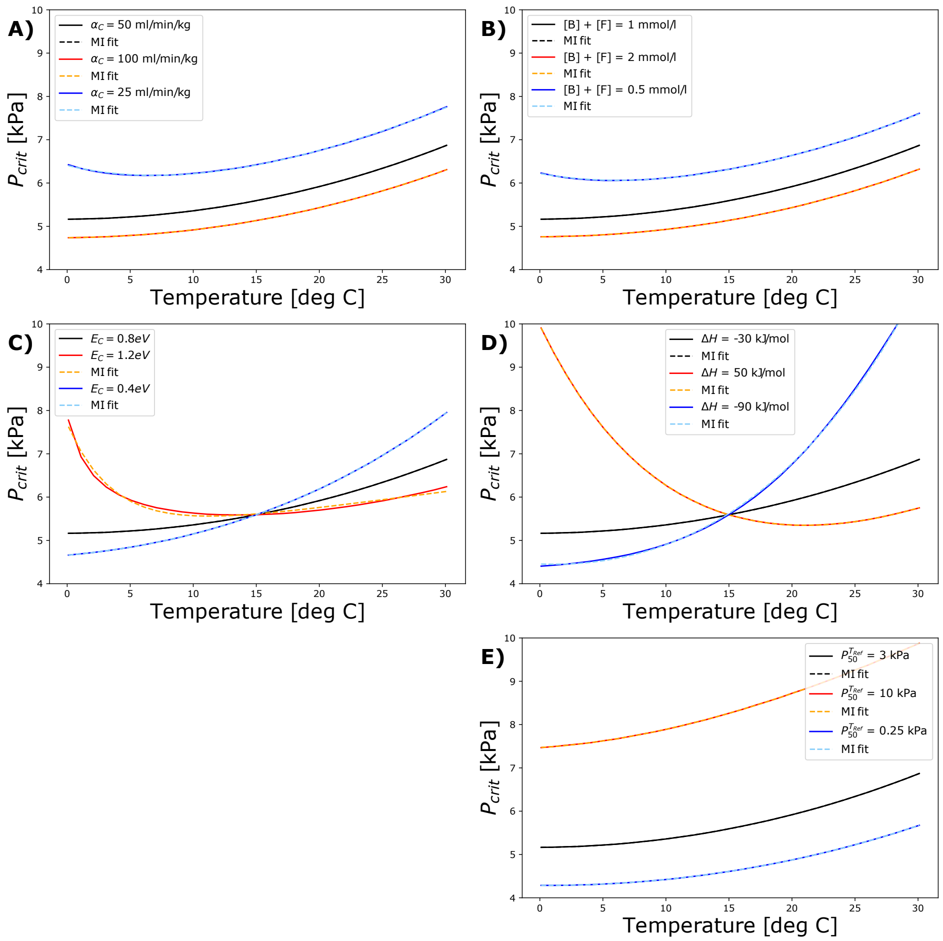


Fig. S4. Variations in the biophysical parameters governing internal circulation (left column) result in similar $P_{crit}$ changes as variations in the chemical parameters describing the transport of $O_2$ by proteins (right column). Fits based on Eqn. (2) from the main text with $n = 2$ supply steps (dashed) capture the simulated $P_{crit}$ curves. **A)** The constant $\alpha_C$ determines the circulation rate at the reference temperature. **B)**. Changes to the total concentration of $O_2$ transport protein have a similar effect as observed for $\alpha_C$. **C)**. The temperature sensitivity of circulation $E_C$ determines the slope of the Arrhenius relationship in SI Eqn. (11). **D)** The enthalpy of the binding reaction changes the effective temperature dependence of the circulation rate similar to changes in $E_C$. **E)** The half-saturation pressure at the reference temperature $P_{50}^{T_{Ref}}$ describes the overall affinity of the transport protein to $O_2$ and correspondingly affects the supply capacity.

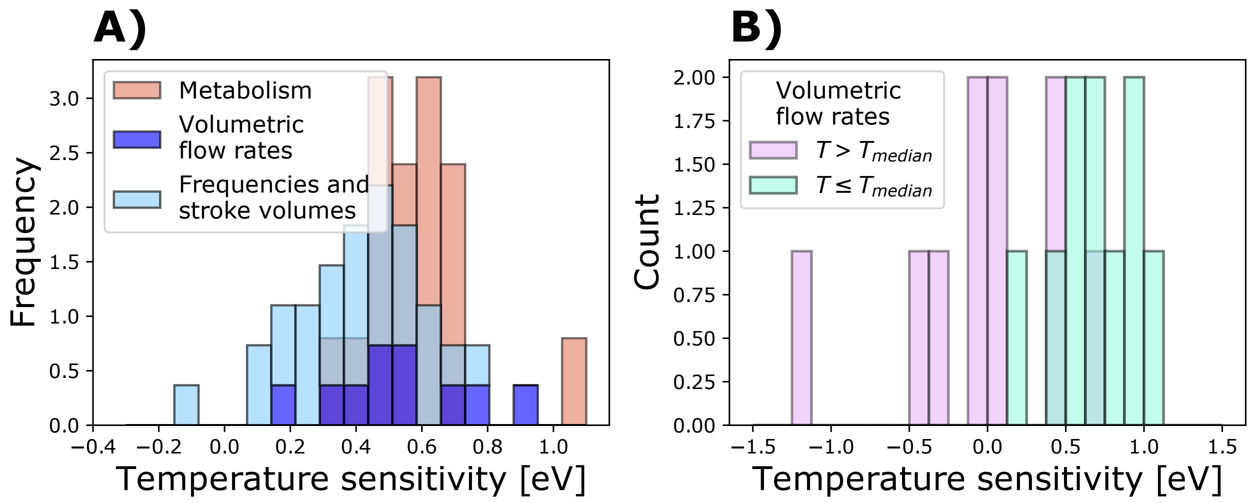


Fig. S5. Frequency distributions of additional estimated temperature sensitivities. **A)** The sensitivity estimates for ventilation and circulation as well as metabolic demand in the 17 species for which both quantities are available yield a distribution that is comparable to that obtained from all species (Fig. 4). **B)** The estimated temperature sensitivity of volumetric ventilation and circulation flow rates below the median experimental temperature far exceed that above the median experimental temperature on average, indicating that these supply processes accelerate at a faster rate under cold conditions.

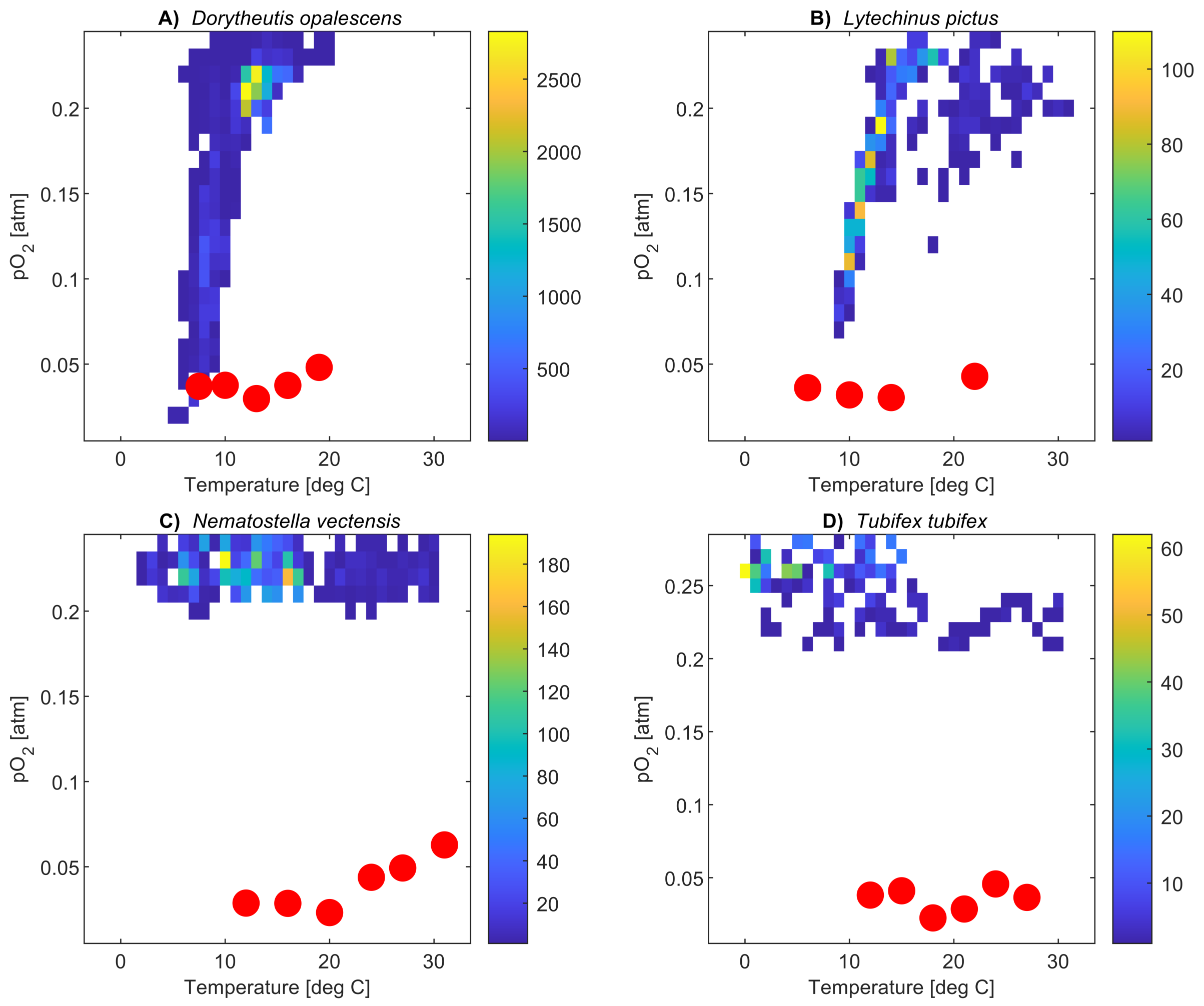


Fig. S6. State-space habitats of OBIS occurrences along with mean experimental $P_{crit}$ values for the 4 species presented in Fig. 1, including **A)** market squid from the California Current System, **B)** an outer shelf sea urchin, **C)** an intertidal anemone and **D)** a sludge worm that mostly inhabits freshwater habitat.

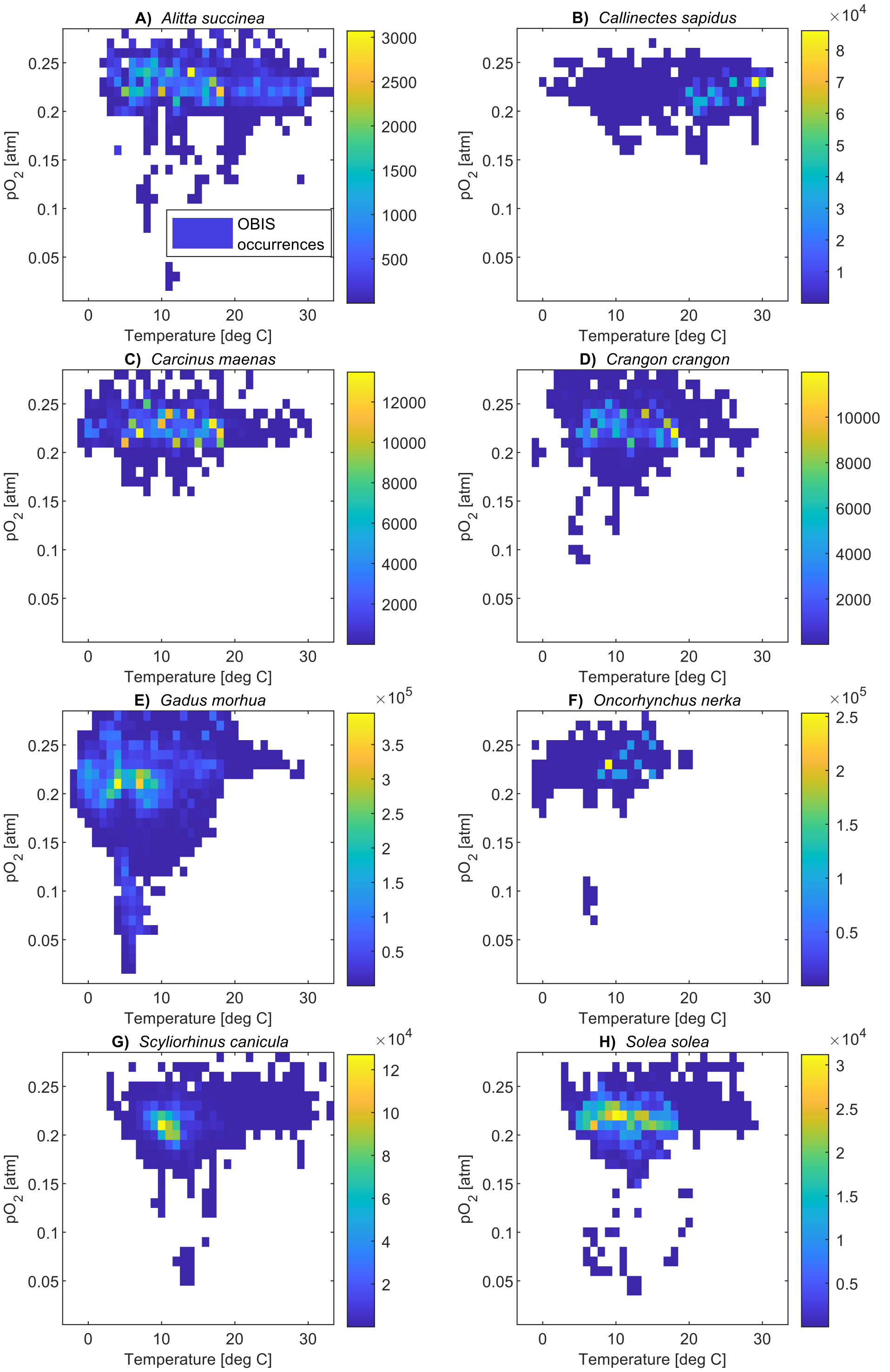


Fig. S7. State-space habitats for all species with sufficient physiological data on temperature-dependent $O_2$ supply and demand rates as well as occurrence data in OBIS.

**Table S1. Descriptions and default values of state variables and parameters in the dynamical model.**

| | Description | Default Value | Unit |
|---|---|---|---|
| $T$ | Temperature | 273.15 – 303.15 | K |
| $k_B$ | Boltzmann's constant | 8.6173e-5 | eV/K |
| $R_{Gas}$ | Gas constant | 0.008314 | kJ/K/mol |
| $T_{ref}$ | Reference temperature | 288.15 | K |
| $\kappa_H(T)$ | Solubility of $O_2$ in seawater | Ref. (7) | $\mu$mol/liter/kPa |
| $\mu$ | Dynamic viscosity of seawater | Ref. (10) | kg/min/s |
| $X_W$ | $O_2$ Concentration in environment | 300 | $\mu$mol/liter |
| $X_B$ | $O_2$ Concentration in boundary layer | 300 | $\mu$mol/liter |
| $X_G$ | $O_2$ Concentration in gill | 300 | $\mu$mol/liter |
| $X_I$ | $O_2$ Concentration in tissue | 300 | $\mu$mol/liter |
| $F_G$ | Free protein concentration in gill | $(1-Y)\cdot 1$ | mmol/liter |
| $F_I$ | Free protein concentration in tissue | $(1-Y)\cdot 1$ | mmol/liter |
| $B_G$ | Bound protein concentration in gill | $Y \cdot 1$ | mmol/liter |
| $B_I$ | Bound protein concentration in tissue | $Y \cdot 1$ | mmol/liter |
| $V_W$ | Volume of environmental compartment | 20 | liter |
| $V_B$ | Volume of boundary layer compartment | 0.005 | liter |
| $V_G$ | Volume of gill compartment | 0.005 | liter |
| $V_I$ | Volume of tissue compartment | 0.005 | liter |
| $\alpha_V$ | Ventilation rate at $T_{ref}$ | 0.002 | liter/min |
| $\alpha_D$ | Diffusion rate at $T_{ref}$ | 0.029 | $\mu$mol /min / kPa |
| $\alpha_C$ | Circulation rate at $T_{ref}$ | 0.00025 | liter/min |
| $\alpha_M$ | (Resting) Metabolic rate at $T_{ref}$ | 0.05 | $\mu$mol/liter/min |
| $E_V$ | Temperature sensitivity of ventilation | 0.8 | eV |
| $E_D$ | Temperature sensitivity of diffusion | 0.21 | eV |
| $E_C$ | Temperature sensitivity of circulation | 0.8 | eV |
| $E_M$ | Temperature sensitivity of metabolism | 0.5 | eV |
| $P_{50}^{T_{ref}}$ | Half-saturation pressure at $T_{ref}$ | 3 | kPa |
| $Y$ | Fraction of bound $O_2$-transport protein | SI Eqn. (15) | - |
| $n_H$ | Hill coefficient | 2 | - |
| $\Delta H$ | Enthalpy of $O_2$ binding reaction | -30 | kJ/mol |
| $K_{eq}$ | Equilibrium constant of $O_2$ binding reaction | $\frac{1}{(P_{50}\kappa_H)^{n_H}}$ | $\mu$mol$^2$ / liter$^2$ |
| $k_f$ | Forward/$O_2$ binding reaction rate | 10 $\alpha_D$ | liter$^3$ / $\mu$mol$^2$ /min |
| $k_b$ | Backward/$O_2$ dissociation reaction rate | $k_f K_{eq}$ | liter/min |

**SI Dataset S1 (separate file, sensitivity_estimates.csv)**

Temperature sensitivity estimates from published data on ventilation and circulation in aquatic water breathers. If available, separate estimates above and below the median experimental temperature as well as estimates of the temperature sensitivity of resting metabolism are included.
