## Supplementary material for "Physiological causes and biogeographic consequences of thermal optima in the hypoxia tolerance of marine ectotherms": SI_Dataset_S1

| Species | Phylum | Habitat | Quantity | Sensitivity [eV] | Cold side [eV] | Warm side [eV] | Reference |
|---|---|---|---|---|---|---|---|
| Acipenser schrenckii | Chordata | Freshwater | vent freq | 0.39 | 0.42 | 0.31 | Zhao ea '17 Aquaculture 48:5338-5345 |
| Acipenser schrenckii | Chordata | Freshwater | vent volume | 0.24 | 0.15 | 0.2 | Zhao ea '17 Aquaculture 48:5338-5345 |
| Acipenser schrenckii | Chordata | Freshwater | vent rate | 0.56 | 0.47 | 0.46 | Zhao ea '17 Aquaculture 48:5338-5345 |
| Acipenser schrenckii | Chordata | Freshwater | metabolic rate | 0.51 | nan | nan | Zhao ea '17 Aquaculture 48:5338-5345 |
| Alitta succinea | Annelida | Marine | vent rate | 0.33 | 0.72 | 0.04 | Kristensen '83 Mar Ecol Prog Ser |
| Alitta succinea | Annelida | Marine | metabolic rate | 0.62 | nan | nan | Cammen '80 MarBiol 61: 9-20 |
| Bidyanus bidyanus | Chordata | Freshwater | vent freq | -0.03 | 0.02 | -0.01 | Patra ea '09 Environ. Toxicol. Chem. 28: 2182-2190 |
| Busycon canaliculatum | Mollusca | Marine | heart rate | 0.75 | nan | nan | Defur & Mangum '79 CBP A 62: 283-294 |
| Callinectes sapidus | Arthropoda | Marine | heart rate | 0.67 | nan | nan | Defur & Mangum '79 CBP A 62: 283-294 |
| Callinectes sapidus | Arthropoda | Marine | metabolic rate | 1.04 | nan | nan | Batterton & Cameron '78 JExpZool 203: 403-418 |
| Cancer magister | Arthropoda | Marine | heart rate | 0.37 | 0.55 | 0.2 | Florey & Kriebel '74 CBP A 48: 285-300 |
| Carcinus maenas | Arthropoda | Marine | vent freq | 0.48 | nan | nan | Taylor ea '73 JCompPhysiol 83: 95-115 |
| Carcinus maenas | Arthropoda | Marine | heart rate | 0.61 | nan | nan | Taylor ea '73 JCompPhysiol 83: 95-115 |
| Carcinus maenas | Arthropoda | Marine | metabolic rate | 0.61 | nan | nan | Klein Breteler '75 JSR 9: 243-254 |
| Channa argus | Chordata | Freshwater | vent freq | 0.35 | 0.49 | 0.33 | Xie ea '17 TURK J FISH AQUAT SC 17: 535-542 |
| Channa argus | Chordata | Freshwater | vent volume | 0.35 | 0.51 | 0.6 | Xie ea '17 TURK J FISH AQUAT SC 17: 535-542 |
| Channa argus | Chordata | Freshwater | metabolic rate | 0.68 | nan | nan | Xie ea '17 TURK J FISH AQUAT SC 17: 535-542 |
| Coryphaena hippurus | Chordata | Marine | vent freq | 0.37 | 0.42 | 0.32 | Szyper & Lutnesky '91 Prog Fish-Cult 53: 166-172 |
| Crangon crangon | Arthropoda | Marine | heart rate | 0.58 | 0.34 | 0.48 | Spaargaren '73 CBP A 45: 773-786 |
| Crangon crangon | Arthropoda | Marine | metabolic rate | 0.47 | nan | nan | Van Donk & Wilde '81 JSR 15: 54-64 |
| Cyprinus carpio | Chordata | Freshwater | heart rate | 0.62 | nan | nan | Stecyk & Farrell '06 Physiol. Biochem. Zool. 79: 614-627 |
| Cyprinus carpio | Chordata | Freshwater | circ volume | 0.12 | nan | nan | Stecyk & Farrell '06 Physiol. Biochem. Zool. 79: 614-627 |
| Cyprinus carpio | Chordata | Freshwater | circulation rate | 0.58 | nan | nan | Stecyk & Farrell '06 Physiol. Biochem. Zool. 79: 614-627 |
| Cyprinus carpio | Chordata | Freshwater | metabolic rate | 0.45 | nan | nan | Clarke & and Johnson, '99 |
| Dentex dentex | Chordata | Marine | vent freq | 0.34 | 0.11 | 0.35 | Valverde et al. '06 |
| Diplodus puntazzo | Chordata | Marine | vent freq | 0.18 | 0.2 | 0.17 | Cerezo & Garcia Garcia '04 J Appl. Ichthyol. 20: 488-492 |
| Gadus morhua | Chordata | Marine | heart rate | 0.26 | 0.56 | 0.11 | Gollock '06 J. Exp. Biol. 209: 2961-2970 |
| Gadus morhua | Chordata | Marine | circ volume | 0.18 | 0.03 | 0.54 | Gollock '06 J. Exp. Biol. 209: 2961-2970 |
| Gadus morhua | Chordata | Marine | circulation rate | 0.4 | 0.61 | 0.65 | Gollock '06 J. Exp. Biol. 209: 2961-2970 |
| Gadus morhua | Chordata | Marine | metabolic rate | 0.67 | nan | nan | Gollock '06 J. Exp. Biol. 209: 2961-2970 |
| Leiopotherapon unicolor | Chordata | Freshwater | vent freq | 0.25 | 0.46 | 0.12 | Gehrke '88 CBP A 89: 587-592 |
| Leiopotherapon unicolor | Chordata | Freshwater | heart rate | 0.39 | 0.67 | 0.27 | Gehrke '88 CBP A 89: 587-592 |

| | | | | | | | |
|---|---|---|---|---|---|---|---|
| Leiopotherapon unicolor | Chordata | Freshwater | metabolic rate | 0.58 | nan | nan | Gehrke '88 CBP A 89: 587-592 |
| Libinia emarginata | Arthropoda | Marine | heart rate | 0.48 | nan | nan | Defur & Mangum '79 CBP A 62: 283-294 |
| Limulus polyphemus | Arthropoda | Marine | heart rate | 0.42 | nan | nan | Defur & Mangum '79 CBP A 62: 283-294 |
| Lysmata seticaudata | Arthropoda | Marine | heart rate | 0.56 | 0.51 | 0.53 | Spaargaren '73 CBP A 45: 773-786 |
| Maja squinado | Arthropoda | Marine | vent freq | 0.04 | 0.7 | -0.68 | Frederich & Portner '00 AM J PHYSIOL-REG I 279: R1531-R1538 |
| Maja squinado | Arthropoda | Marine | heart rate | 0.11 | 0.45 | -0.33 | Frederich & Portner '00 AM J PHYSIOL-REG I 279: R1531-R1538 |
| Melanotaenia duboulayi | Chordata | Freshwater | vent freq | 0.11 | 0.16 | 0.22 | Patra ea '09 Environ. Toxicol. Chem. 28: 2182-2190 |
| Nereis diversicolor | Annelida | Marine | vent rate | 0.22 | 0.71 | -0.06 | Kristensen '83 Mar Ecol Prog Ser |
| Nereis virens | Annelida | Marine | heart rate | 0.54 | 0.88 | -0.01 | Defur & Mangum '79 CBP A 62: 283-294 |
| Nereis virens | Annelida | Marine | vent rate | 0.3 | nan | nan | Kristensen '83 Mar Ecol Prog Ser |
| Noetia ponderosa | Mollusca | Marine | heart rate | 0.9 | nan | nan | Defur & Mangum '79 CBP A 62: 283-294 |
| Oncorhynchus mykiss | Chordata | Anadromous | vent freq | 0.09 | 0.24 | -0.16 | Patra ea '09 Environ. Toxicol. Chem. 28: 2182-2190 |
| Oncorhynchus nerka | Chordata | Anadromous | circulation rate | 0.18 | 0.24 | 0.06 | Davis '68 PhD diss. UBC |
| Oncorhynchus nerka | Chordata | Anadromous | circ volume | -0.14 | -0.14 | -0.22 | Davis '68 PhD diss. UBC |
| Oncorhynchus nerka | Chordata | Anadromous | heart rate | 0.46 | nan | nan | Davis '68 PhD diss. UBC |
| Oncorhynchus nerka | Chordata | Anadromous | metabolic rate | 0.47 | nan | nan | Davis '68 PhD diss. UBC |
| Oncorhynchus nerka | Chordata | Anadromous | vent freq | 0.46 | nan | nan | Davis '68 PhD diss. UBC |
| Orthodon microlepidotus | Chordata | Freshwater | vent freq | 0.17 | nan | nan | Campagna & Cech '81 J. Fish Biol. 19: 581-591 |
| Orthodon microlepidotus | Chordata | Freshwater | vent volume | 0.4 | nan | nan | Campagna & Cech '81 J. Fish Biol. 19: 581-591 |
| Orthodon microlepidotus | Chordata | Freshwater | vent rate | 0.67 | nan | nan | Campagna & Cech '81 J. Fish Biol. 19: 581-591 |
| Orthodon microlepidotus | Chordata | Freshwater | metabolic rate | 0.48 | nan | nan | Campagna & Cech '81 J. Fish Biol. 19: 581-591 |
| Pagothenia borchgrevinki | Chordata | Marine | vent freq | 0.43 | 0.24 | 0.8 | Robinson ea '11 Polar Biology 34: 371-379 |
| Pagothenia borchgrevinki | Chordata | Marine | heart rate | 0.25 | 0.15 | 0.44 | Robinson ea '11 Polar Biology 34: 371-379 |
| Palaemon serratus | Arthropoda | Marine | heart rate | 0.14 | 0.17 | 0.14 | Spaargaren '73 CBP A 45: 773-786 |
| Pholis gunnellus | Chordata | Marine | vent freq | 0.23 | 0.28 | 0.22 | Laming '83 CBP A 76: 71-73 |
| Piaractus mesopotamicus | Chordata | Freshwater | vent freq | 0.42 | 0.55 | 0.26 | Aguiar ea '02 J. Therm. Biol. 27:299-308 |
| Piaractus mesopotamicus | Chordata | Freshwater | vent volume | 0.08 | 0.03 | 0.16 | Aguiar ea '02 J. Therm. Biol. 27:299-308 |
| Piaractus mesopotamicus | Chordata | Freshwater | vent rate | 0.5 | 0.58 | 0.45 | Aguiar ea '02 J. Therm. Biol. 27:299-308 |
| Piaractus mesopotamicus | Chordata | Freshwater | heart rate | 0.54 | 0.72 | 0.26 | Aguiar ea '02 J. Therm. Biol. 27:299-308 |
| Piaractus mesopotamicus | Chordata | Freshwater | metabolic rate | 0.35 | nan | nan | Aguiar ea '02 J. Therm. Biol. 27:299-308 |
| Platichthys stellatus | Chordata | Marine | vent rate | 0.9 | 0.89 | -1.21 | Watters & Smith '73 Marine Biology 19: 133-148 |
| Platichthys stellatus | Chordata | Marine | circulation rate | 0.51 | 0.74 | 0.27 | Watters & Smith '73 Marine Biology 19: 133-148 |
| Platichthys stellatus | Chordata | Marine | metabolic rate | 0.68 | nan | nan | Watters & Smith '73 Marine Biology 19: 133-148 |

| | | | | | | | |
|---|---|---|---|---|---|---|---|
| Prochilodus scrofa | Chordata | Freshwater | vent freq | 0.53 | -0.05 | 1.02 | Fernandes ea '95 J. Fish Biol. 46: 123-133 |
| Prochilodus scrofa | Chordata | Freshwater | vent volume | 0.29 | 0.94 | -0.38 | Fernandes ea '95 J. Fish Biol. 46: 123-133 |
| Prochilodus scrofa | Chordata | Freshwater | vent rate | 0.77 | 0.76 | 0.63 | Fernandes ea '95 J. Fish Biol. 46: 123-133 |
| Prochilodus scrofa | Chordata | Freshwater | metabolic rate | 0.58 | nan | nan | Fernandes ea '95 J. Fish Biol. 46: 123-133 |
| Sander lucioperca | Chordata | Freshwater | vent freq | 0.75 | 0.59 | 1.11 | Frisk ea '12 Aquaculture 324:151-157 |
| Sander lucioperca | Chordata | Freshwater | metabolic rate | 0.56 | nan | nan | Frisk ea '12 Aquaculture 324:151-157 |
| Scyliorhinus canicula | Chordata | Marine | heart rate | 0.46 | nan | nan | Taylor ea '77 J. Exp. Biol. 70: 57-75 |
| Scyliorhinus canicula | Chordata | Marine | metabolic rate | 0.64 | nan | nan | Butler and Taylor, '75 |
| Solea solea | Chordata | Marine | metabolic rate | 0.41 | nan | nan | Lefrancois & Claireaux '03 Mar. Ecol. Prog. Ser 259: 273-284 |
| Solea solea | Chordata | Marine | heart rate | 0.63 | 0.55 | 0.7 | Lefrancois & Claireaux '03 Mar. Ecol. Prog. Ser 259: 273-284 |
